# Spatial transcriptomics identifies dysregulated programs across neural and non-neural tissues in spinal muscular atrophy

**DOI:** 10.64898/2026.03.20.713154

**Authors:** Anne Rietz, Lokesh Kumari, Elliot J. Androphy

## Abstract

Spinal muscular atrophy (SMA) is caused by insufficient levels of the survival motor neuron (SMN) protein and clinically manifests as profound weakness due to motor neuron degeneration. While recent evidence suggests it is a multisystem disorder, the pathological programs in neural and peripheral tissues are poorly understood. We applied spatial transcriptomics to cross-sections from the entire lumbar region of pre-symptomatic SMA and control mice to define early, tissue-wide transcriptional consequences of SMN deficiency. This approach enabled spatially preserved analysis of spinal cord, dorsal root ganglia, muscle, bone, cartilage, bone marrow, adipose, and connective tissues within their native anatomical context. Within the SMA spinal cord, motor neuron associated regions and ventral interneurons exhibited upregulation of neurofilaments, tubulin isoforms, and microtubule transport machinery. Neurofilament changes were largely restricted to motor neurons, and tubulin dysregulation extended broadly across ventral regions.

Multiple tissues displayed collagen and extracellular matrix gene dysregulation at post-natal day 4. Skeletal muscle demonstrated fiber type-specific stress responses, including induction of atrophy-associated genes, while bone displayed an osteoclast-dominant transcriptional signature, consistent with accelerated resorption and a pro-osteoporotic state. Bone marrow transcriptomes indicated activation of neutrophil degranulation, innate immune signaling, and osteoclastogenic pathways, identifying bone marrow as an active inflammatory and skeletal regulatory niche in SMA. Adipose tissue exhibited extracellular matrix dysregulation, complement activation, profibrotic TGF-β signaling, and stress-induced lipolysis.

These findings reveal that SMN deficiency drives early transcriptional reprogramming across multiple tissues well before motor neuron loss and identify non-neuronal pathological programs that offer therapeutic targets for improving long-term outcomes in SMA.

## INTRODUCTION

Spinal muscular atrophy (SMA) is an inherited autosomal recessive neurodegenerative disease that leads to loss of motor neurons. The pathogenesis of SMA is linked to the survival motor neuron gene SMN. The human genome normally encodes two SMN alleles called SMN1 and SMN2, which have identical coding sequence. Absence of the SMN1 genes is partially compensated by SMN2. Due to a single exonic nucleotide difference from SMN1, the majority of SMN2 transcripts lack exon 7 1,2. The low level of SMN protein produced from the full length SMN2 mRNA is sufficient for embryonic development, however, as the infant grows, progressive motor weakness is typically fatal. Some individuals have multiple copies of SMN2 and develop milder symptoms at later timepoints. Two marketed therapeutics enhance inclusion of exon 7 in the SMN2 transcript and an SMN1 transgene viral vector enable partial recovery of motor skills. SMN is ubiquitously expressed, and extended survival of patients receiving SMN-restoring therapy has revealed extra-neuronal pathology 3.

Mice encode a single homologue of human *SMN*1 (*mSmn*). While mice heterozygous with a single allele do not display any pathology, *mSmn* null is early embryonic lethal. Transgenic mouse models lack *mSmn* and carry one or more human *SMN2* transgenes, which make sufficient SMN protein for embryonic development. These animals are small, become weak, and die in 10-14 days. We performed spatial transcriptomics (ST) experiments using SMA mice to quantitatively evaluate and compare the transcriptomes of post-natal day 4 (PND4) normal and pre-symptomatic mice. By using whole cross sections of the lumbar region, analysis of the entire spinal motor and sensory regions as well as flanking muscle, bone, cartilage, marrow, and adipose were characterized.

## RESULTS

Lumbar (L4–L5) cross-sections were collected from four female heterozygous control and four SMA mice from two independent litters at PND4. Sections were H&E stained and showed spinal cord, dorsal root ganglia, muscle, adipose and non-decalcified bone (Figure scol1A). RNA integrity of FFPE samples exceeded Visium requirements (DV200 >93%; Figure S1A). Sequencing and Space Ranger processing generated an average of 119 million reads per chip (∼475 million total), with transcripts detected from ∼19,291 genes and 89% mapping confidence (Table S1). Slightly fewer spatial transcriptomic spots were captured in SMA samples compared to controls (4505 vs. 5833), consistent with reduced body size. Spatial mapping of ST spots is shown in Figure 1B.

**Figure 1.**
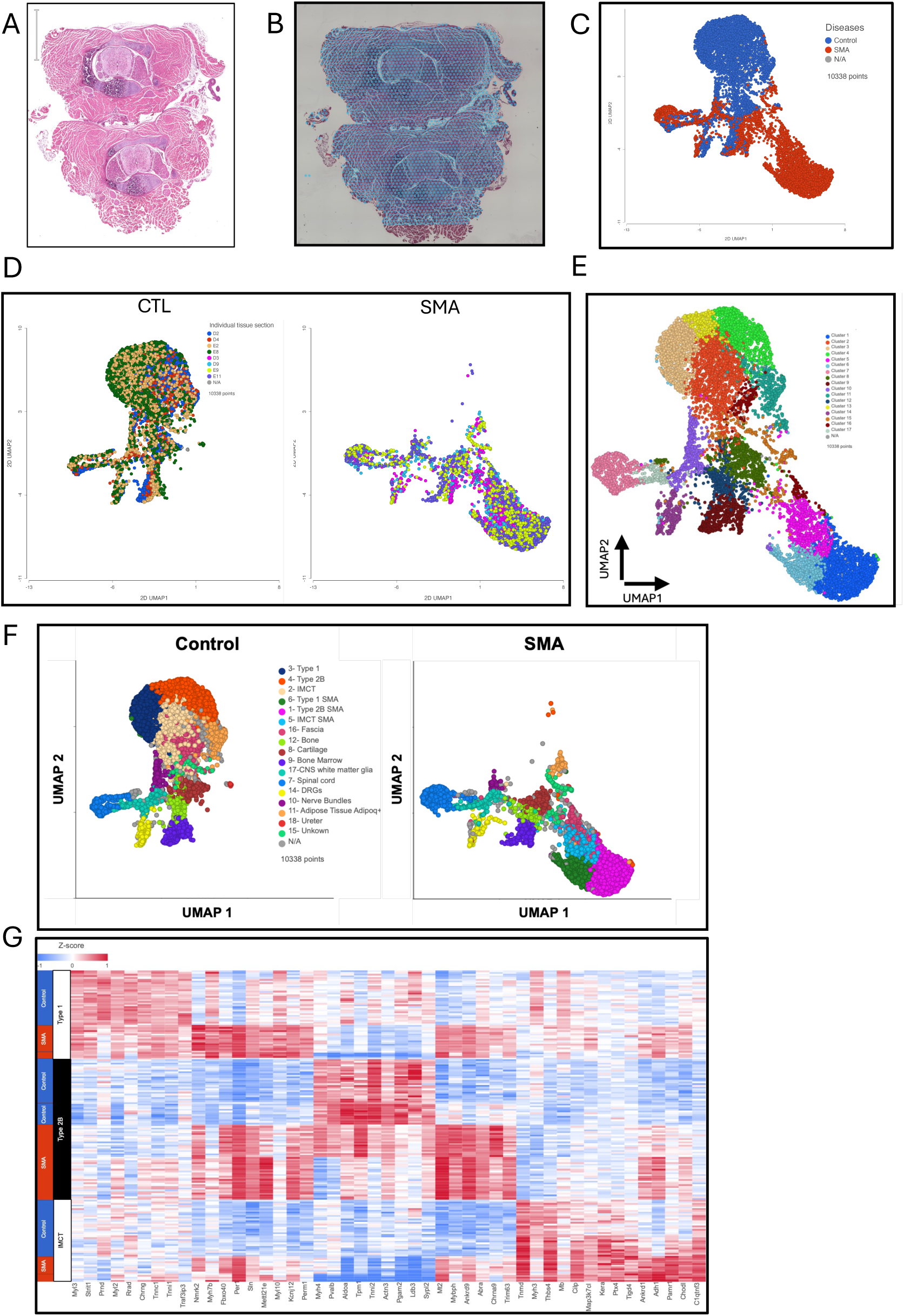
Spatial transcriptomics (ST) and histology identify multiple tissue types. **(A)** H&E stain and **(B)** ST spots of two L4–L5 lumbar spinal columns cross sections from control mice. Each hexagon represents a Visium RNA capture point generated in Partek Flow.G **(C)** UMAP projection from all animals colored by genotype (control-blue; SMA-red). **(D)** UMAP plot showing ST spots from four control and four SMA biological replicates. **(E)** Graph-based unsupervised clustering of ST spots identifying 17 transcriptionally distinct clusters across all tissues. **(F)** UMAP plots displaying cluster annotations for control and SMA animals based on biomarker gene expression and anatomical localization. **(G)** Heatmap of biomarker expression in cluster-defining genes across annotated muscle tissues and genotypes.

A uniform manifold approximation and projection (UMAP) visualization of unsupervised clustering generated in Space Ranger revealed distinct RNA expression patterns across ST spots (Figure 1E). Coloring spots by condition (control, blue; SMA, red) highlighted clear differences between groups (Figure 1C). Reproducibility across samples was confirmed by clustering of individual spinal columns within each cohort (Figure 1D). Graph-based clustering analysis identified 17 distinct clusters (Figure 1E). Several clusters were shared between control and SMA samples, although clear differences were observed (Figure 1F). Shared clusters were annotated based on marker gene expression and anatomy, corresponding to spinal cord, dorsal root ganglion (DRG), bone, cartilage, bone marrow, fascia and adipose (Figure 1F, Figure S1D). For example, cluster 16 represents posterior deep fascia, enriched for extracellular matrix (ECM) genes (*Egfl6*, *Kera*, *Ecrg4, C1qtnf3*, *Cpxm2*, *Ssc5d*, *Sfrp2*, *Gpnmb*, *Abi3bp*, *Zfp185)* and tendon-associated genes (*Tnmd*, *Mkx*, *Fmod*, *Sparc, Thbs4*).

### Pathological alterations in muscle

Clusters 2, 3, 4, and 13 were absent in SMA, while clusters 1, 5, 6 were SMA-specific and identified as muscle by marker expression and histology. Cluster 6 corresponded to Type 1 muscle (*Myl3*, *Myl2, Tnni1, Tnnc1, Tnnt1*), whereas clusters 4 and 13 (merged as cluster 4) represented Type 2b muscle (*Myh4*, *Aldoa*, *Pvalb*, *Actn3*, *Tnni2*; Figure 1E vs. 1F). These assignments were supported by spatial heatmaps and low *Myh2* expression in Type 2b regions (Figures S2A–C). Cluster 2 aligned with intramuscular connective tissue (IMCT) based on fibroblast/tendon biomarker gene expression (*Tnmd, Thbs4, Cilp, Kera, Ptx4, Tigd4, Map3k7cl*). Type 1, Type 2b, and IMCT clusters present in controls were replaced by SMA-specific clusters 1, 5, and 6, which retained fiber-type identity but exhibited shared stress-response signatures, including circadian/metabolic regulation (*Per1*), atrophy-associated pathways (*Mettl21e*, *Fbxo40*, *Trim63*), and calcium–energy coupling (*Sln*; Figure 2A, Figure S2D).

**Figure 2.**
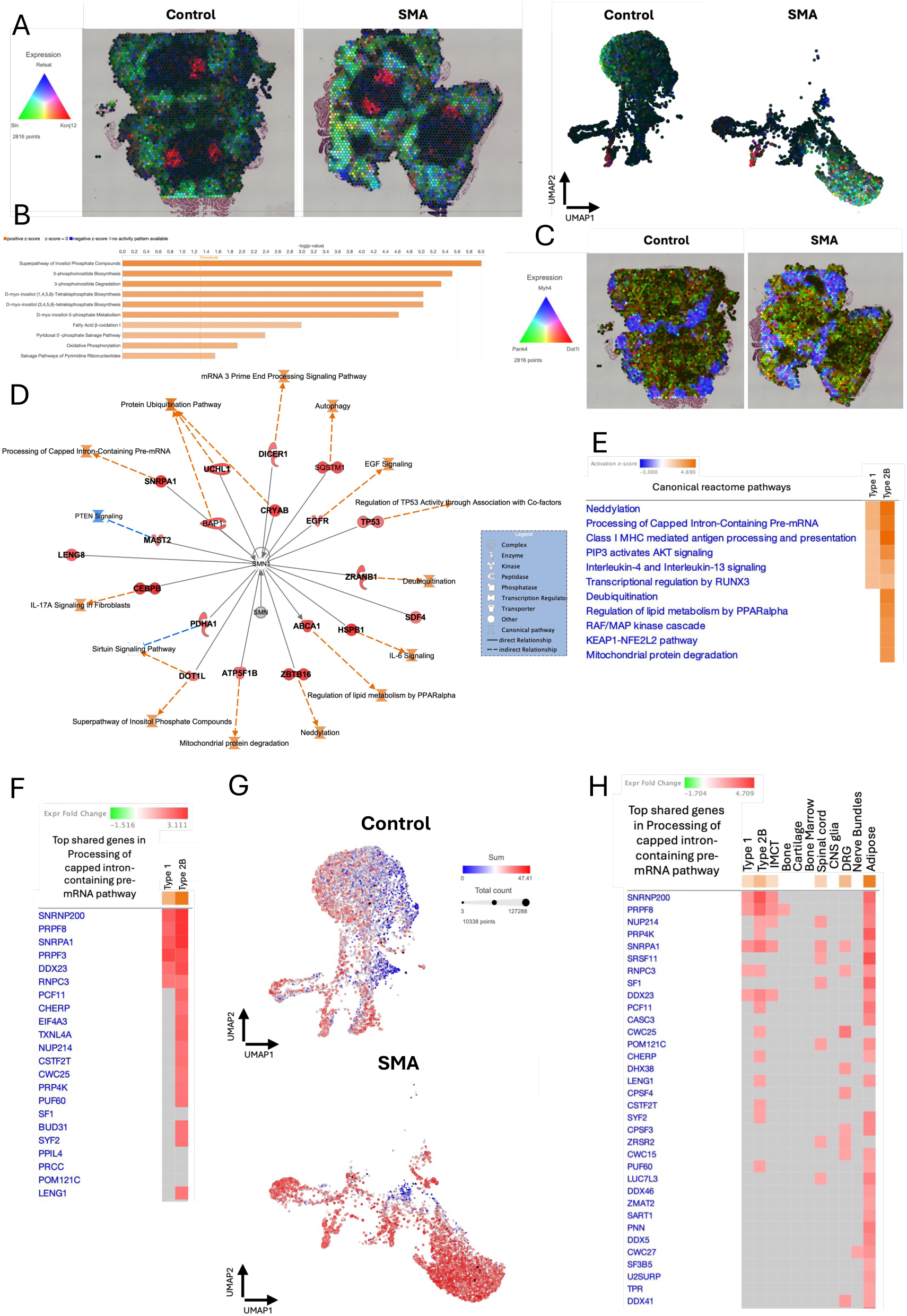
ST analysis identifies SMA-specific muscle clusters and dysregulation of RNA processing, metabolic, and stress-response pathways. **A)** Spatial heatmaps (left) and UMAP projections (right) showing expression of the top biomarkers defining the SMA-specific Type 2b muscle cluster (Retsat-blue, Sln-green, Kcnj12-red). Color intensity correlates with transcript abundance and white spots indicate co-expression of all three genes. **B)** Ingenuity metabolic pathway analysis (IPA) of differentially expressed genes in SMA Type 2b muscle cluster with a z-score ≥2 considered significant. A positive z-score (orange) indicates pathways activation, and negative-score (blue) indicates inhibition. **C)** Spatial expression heatmap of representative metabolic and signaling genes (*Pank4*-green, *Dot1l*-red) and *Myh4* (blue) indicate Type 2b fibers in control and SMA sections. **D)** SMN-centered interaction network using Qiagen IPA PathDesigner. Identification of differentially expressed genes (DEG) associated with SMA Type 2b muscle connected to SMN through protein-protein or protein-RNA interactions. **E)** IPA Reactome comparison pathway analysis of DEGs in Type 1 and Type 2b SMA muscle (z-score ≥2). **F)** Gene expression heatmap of the top shared genes involved in the processing of capped intron-containing pre-mRNA pathway in Type 1 and Type 2b SMA muscle. **G)** Projection of pre-mRNA processing pathway genes (SUM) onto the spatial UMAP representation. Red intensity indicates higher transcript expression and dot size indicates total transcript count in the ST spot. **H)** Cross-tissue comparison of the top genes involved in the processing of capped intron-containing pre-mRNA pathway across multiple SMA tissues, including Type 1 and Type 2b muscle, bone marrow, cartilage, spinal cord, dorsal root ganglia (DRG), nerve bundles, and adipose tissue.

Top biomarkers in Type 1 SMA muscle indicate activation of pathways that mediate i) adaptive slow-oxidative mitochondrial and redox stress (*Nmrk2, Perm1, Retsat*); ii) Ca²⁺ handling stress and energy inefficiency via *Sln*-mediated SERCA activity ^4,5^; and iii) changes associated with damage or atrophy indicated by increased *Fbxo40* and *Mettl21e* (Tables S2, S3). *Fbxo40* can promote catabolic atrophy via inhibition of the IGF1/PI3K/Akt signaling pathway^6^. Together, these data imply Type 1 SMA muscle initiates atrophy by PND4 and is accompanied by compensatory upregulation of oxidative, endurance, and stress resistance pathways.

Biomarkers in Type 2b SMA muscle indicate fast-glycolytic fibers undergoing degenerative stress (Table S3). Specifically, *Trim63* promotes atrophy, while increased *Mt2, Ankrd9,* and *Per1* expression likely reflect metabolic and oxidative adaptation. Concurrent upregulation *of Abra, Mybph,* and *Mettl21e* suggest compensatory hypertrophic and structural processes to maintain contractile function^7^. DESeq2 analysis further identified atrophy-associated differentially expressed genes (DEGs; *Fbxo32*, *Foxo1/3*; Table S4)^8^. Metabolic dysregulation was also evident including altered inositol phosphate signaling (e.g. *Pank4, Dot1l, Pik3ca;* Figure 2B, C) and fatty acid β-oxidation (e.g. *Acaa2, Acadm, Eci2*). Ingenuity Pathway Analysis (IPA) reactome analysis highlighted enrichment of pathways related to neddylation, lipid metabolism by PPARα, circadian regulation, and interleukin-4 and -13 (Figure S2E, F).

We used Qiagen’s IPA PathDesigner to examine potential relationships between SMN and the 577 DEGs identified in SMA Type 2b muscle. This identified 19 DEGs with reported direct protein–protein or protein–RNA interactions with SMN (Figure 2D). PathDesigner further connected these genes through activation or inhibition relationships to the most dysregulated pathways in Type 2b SMA muscle. Two DEGs, *Leng8* ^9^, which encodes a key RNA nuclear retention factor, and *Sdf4* ^10^, which encodes a Golgi-ER calcium binding protein, were not connected to an altered pathway in SMA Type 2b muscle. Several established SMN dependent proteins were identified such as UCHL1, SQSTM1 *(p62*), p53, and the U2 snRNP SNRPA1 ^11–16^. Neddylation emerged as a top dysregulated pathway potentially linked to SMN via *Zbtb16* (PLZF), a transcriptional repressor that can be part of the CRL3 E3 ubiquitin ligase complex and which increased >3-fold in SMA muscle (Figure S2G).

Type 2b SMA muscle exhibited more DEGs than Type 1 (577 vs. 112; Table S5), with 63 shared DEGs. Both fiber types showed induction of stress and atrophy-associated genes *Ddit4, Gadd45a, Gadd45g, Cdkn1a, Foxo3, Trp53inp1,* and *Phlda3*, which are linked to mTOR downregulation, proteolysis, autophagy, and cell cycle arrest, metabolic and redox signaling (*Abca1, Acot2, Pnpla2, Retsat, Mt1, Mt2, Pex6,* and *Gramd1b*), alongside altered inositol phosphate signaling (*Pik3r1, Pik3c3, Ppp2r1a*, and *Ppp1r3a*; Figure S2H-J).

With regards to pathways directly related to SMN function, pre-mRNA processing (e.g. *Ddx23, Rnpc3, Snrnp200, Snrpa1, Prpf3, Prpf8*) and neddylation were enriched^17,18^ (Figure 2E,F). Projection of shared DEGs within the pre-mRNA regulatory pathway onto the UMAP revealed enrichment of components across multiple tissues (Figure 2G). Cross-tissue analyses confirmed modulation of this pathway in IMCT, spinal cord, DRG, and adipose; with *Snrpa1* as the only consistently shared DEG (Figure 2H; Figure S3). Snrpa1 was reported to be dysregulated across multiple SMA-affected tissues^15^.

### Pathological alterations in bone and bone marrow

ST enabled identification of changes in bone that may be relevant to scoliosis and fractures, which are clinically refractory complications of SMA ^19–26^. SMA bone exhibited 93 DEGs (Table S6) with enrichment of osteoclast-related pathways, including ‘Role of Osteoclasts in Rheumatoid Arthritis’ signaling (Figure 3A). Key osteoclast genes *Spp1* (Oste pontin), *Acp5* (Tartrate-resistant acid phosphatase), and *Ctsk* (Cathepsin K) were increased >2-fold, indicating increased resorptive activity (Figure 3B-D), while their levels were unchanged in SMA cartilage. Elevated expression of matrix metalloproteinases (*Mmp*9, *Mmp13*; >1.5-fold; FDR < 0.05; Figure 3E) and proteases *Ctsb*, *Ctsd,* and the extracellular matrix protease regulator *Serpine2* further support a protease-driven program targeting collagen and proteoglycans (>1.5-fold; FDR < 0.05; Figure 3E). Although cell numbers cannot be directly assessed, the preferential induction of resorption and protease genes suggests enhanced osteoclast function rather than increased abundance. Upregulation of *Cst3* (cystatin C; ∼2 fold; Figure 3E), indicates anti-protease feedback, while increased *Fst* (follistatin; 3-fold; Figure 3F), a regulator of activin/TGF-β signaling and osteoblast–osteoclast coupling, suggests compensatory paracrine responses.

**Figure 3.**
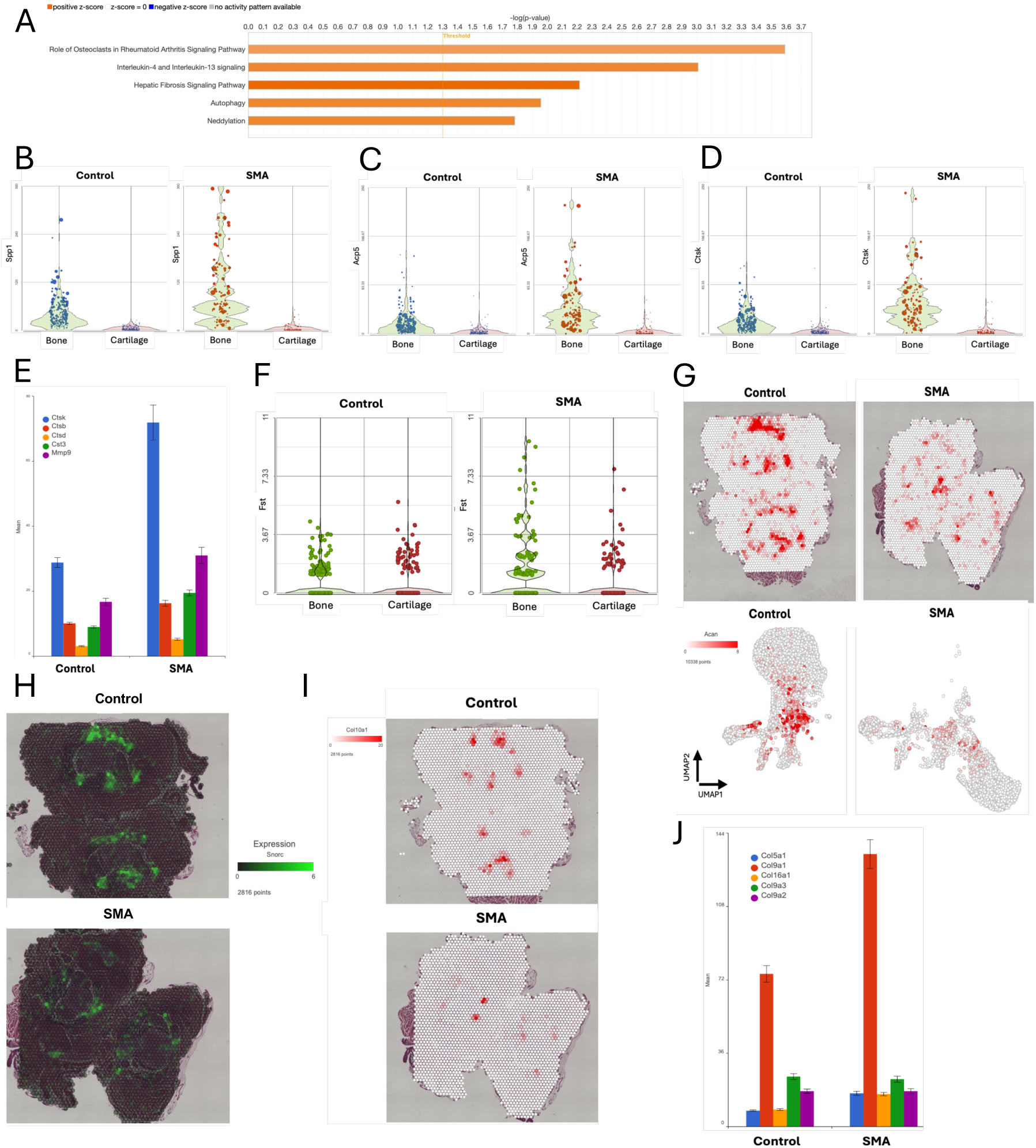
Enhanced osteoclast activity and extracellular matrix remodeling in SMA bone with concurrent disruption of cartilage matrix programs. **A)** IPA of DEGs (FC > 2; FDR < 0.05) identified the most significantly enriched pathways (z-score > 2) in SMA bone. **B-D**) Violin plots showing ST levels of osteoclast-associated genes *Spp1* (B), *Acp5* (C), and *Ctsk* (D) in bone and cartilage from control and SMA. **E)** Bar plots show DESeq2 normalized mean counts of protease and matrix remodeling gene expression in bone. Increased expression of *Ctsk, Ctsb, Ctsd*, and *Mmp9* indicates activation of a protease-driven extracellular matrix degradation program. Upregulation of *Cst3* (cystatin C) suggests engagement of compensatory anti-protease feedback response. **F)** Violin plot showing raw UMI expression of *Fst* (follistatin) in ST spots from bone and cartilage clusters. **G)** Representative spatial heat maps (top) and UMAPs (bottom) showing Acan expression. **H)** Spatial distribution of *Snorc* (green), a marker of differentiated chondrocytes and cartilage matrix homeostasis, demonstrating reduced expression in SMA. **I)** Spatial expression of Col10a1 (red) in cartilage. **J)** Bar plots show DESeq2 normalized mean counts of major structural matrix components *Col10a1* and *Col1a1* and minor fibrillar collagens *Col9a1, Col5a1*, and *Col16a1* in cartilage.

Only 36 genes were differentially expressed in SMA cartilage (Table S7), yet key structural and lineage pathways were disrupted. Expression of *Acan* and *Snorc*, markers of cartilage matrix integrity and chondrocyte maturation, was significantly reduced (Figure 3G, H). Consistent with this, major cartilage collagens (*Col10a1*, *Col1a1*) were decreased while minor fibrillar collagens (*Col9a1, Col5a1, Col16a1*) were increased (Figure 3I,J), suggesting extracellular matrix reprogramming toward stress-associated remodeling rather than normal maturation^27^.

ST revealed pronounced inflammatory activation in SMA bone marrow, exemplified by increased neutrophil and inflammatory monocyte signatures (*S100a8*, *S100a9*) and enrichment of neutrophil degranulation and pathogen-induced cytokine storm pathways (Figure 4A,B, Table S8). This was accompanied by elevated acute-phase and inflammatory mediators *Ptx3, Chil1, Hp,* chemokines *Ccl6* and *Ccl9*, and myeloid inhibitory receptors *Cd33* and *Pirb*, suggesting dysregulated inflammatory control (Figure 4C). Consistent with this, neutrophil lineage and maturation markers *Ngp, Retnlg, Lcn2, Olfm4, Csf3r* were upregulated, implying enhanced production and mobilization. Concurrent induction of neutrophil effector and degranulation genes *Pglyrp1, Stfa3, Ncf1, Alox5ap, Stxbp2* supports a primed inflammatory state. Increased *Mmp8* expression links neutrophil activation to matrix modification, providing a mechanistic connection to the pro-resorptive skeletal phenotype in SMA (Figure 4E).

**Figure 4.**
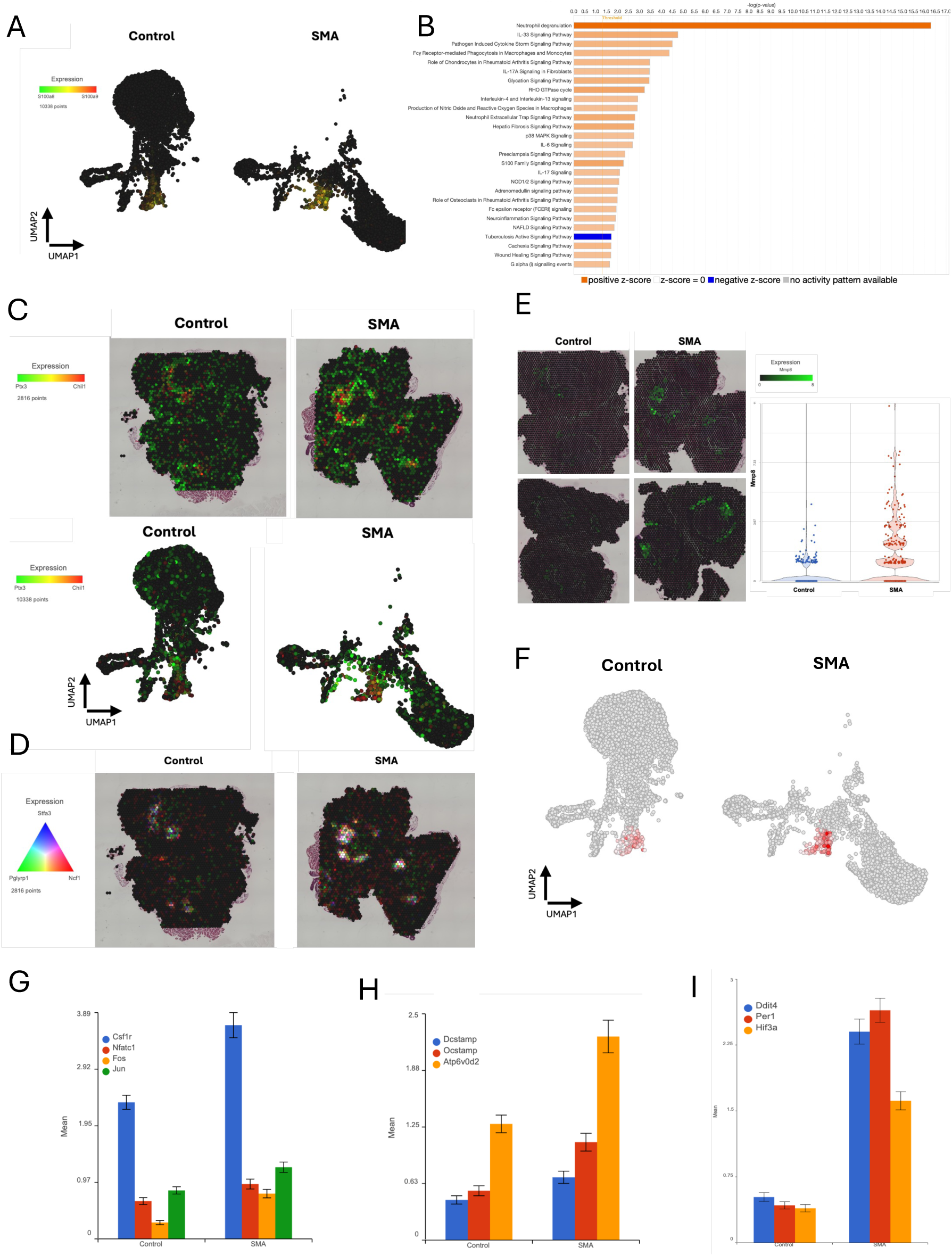
Altered inflammatory and osteoclast-associated programs in SMA bone marrow. **A)** UMAP of ST spots colored by expression of neutrophil and inflammatory monocyte markers *S100a8* (green) and *S100a9* (red), demonstrating increased co-expression (yellow) in SMA bone marrow. **B)** IPA) of DEGs in SMA bone marrow. Positive z-score (orange) indicates pathway activation; negative z-score (blue) indicates inhibition. **C)** Spatial expression of acute-phase and inflammatory mediators *Ptx3* (green) and *Chil1* (red) in bone marrow (top) and UMAP plots (bottom). **D)** Spatial heatmap of neutrophil lineage and effector genes *Pglyrp1* (green), *Stfa3* (blue), and *Ncf1* (red) s. White ST spots indicate co-expression of all three genes. **E)** Spatial heatmap of *Mmp8* expression encoding neutrophil collagenase (green, raw UMI). Violin plots UMI counts of *Mmp8* in bone marrow. Increased color intensities represent increasing UMI counts and yellow represents co-expression. **F)** UMAP plot highlighting spots enriched for the RNA-binding protein *Zfp36l2* (red), a regulator of inflammatory cytokine mRNA stability, in bone marrow cluster. Increased red intensity correlates to increased UMI counts. **G-I)** Bar plots show DESeq2 normalized mean counts for osteoclastogenic and stress-response gene programs in bone marrow. **G)** Increased expression of osteoclast precursor and differentiation markers *Csf1r, Nfatc1, Fos, Jun*. **H)** Upregulation of osteoclast fusion and maturation genes *Dcstamp, Ocstamp, Atp6v0d2*. **I)** Increased expression of circadian and metabolic stress regulators *Ddit4, Per1, Hif3a*. Data presented as mean expression with error bars indicating S.E.M.

This inflammatory state was accompanied by activation of cellular stress and endocrine feedback pathways, including oxidative/ER stress-responsive genes *Mt1, Mt2, Ddit3, Ddit4* and glucocorticoid-regulated targets *Tsc22d3 and Fkbp5*. Upregulation of adhesion and migration regulators *Vcam1, Dock2, Vav1, Arhgap9* suggests enhanced myeloid cells egress and tissue infiltration. Upstream analysis predicts lipopolysaccharide (LPS) as the most significant regulator, followed by TNF, IL1B, and TCL1A (Table S9), revealing a dominant neutrophil- and monocyte mediated inflammatory program in SMA bone marrow. Increased expression of the RNA-binding protein *Zfp36l2*, a key post-transcriptional regulator of inflammatory cytokine mRNA stability, indicates engagement of compensatory post-transcriptional feedback program to restrain cytokine output under chronic inflammatory stress (Figure 4F). Concurrently, the SMA bone marrow exhibits a pro-osteoclastogenic signature with induction of osteoclast specification (*Csf1r, Nfatc1, Fos, Jun*; FDR<0.05; FC≥1.4; Figure 4G), fusion/maturation (*Dcstamp*, *Ocstamp*, *Atp6v0d2*; Figure 4H), and resorption machinery (*Acp5*, *Ctsk*, *Mmp9*), alongside elevated *Tnfsf11* (RANKL) and myeloid markers (*Itgam*). These findings indicate a bone marrow microenvironment that promotes osteoclast activation and bone resorption in SMA, further accompanied by upregulation of circadian and metabolic stress regulators (*Per1*, *Hif3a*, *Ddit4*; Figure 4I).

### Pathological alteration of adipose and connective tissue

DESeq2 analysis of adipose ST indicates that SMA white adipose undergoes fibrotic–inflammatory alterations with stress-driven lipolytic activation rather than adaptive thermogenic pathway induction (Table S10). ECM and basement membrane genes *Col6a1*, *Lama4*, *Nid1*, and *Prelp* were upregulated and spatially co-enriched within adipose microenvironments (Figure 5A,B; Figure S4A-C). Concurrent induction of profibrotic pathways including TGFβ-SMAD *(Tgfb1, Tgfbr3, Smad1, Smad4)* and PDGF signaling (*Fgfr1, Pdgfra*), along with *Igfbp7* and *Igf2bp2* supports active ECM expansion and fibrosis (Figure 5C,D).

**Figure 5.**
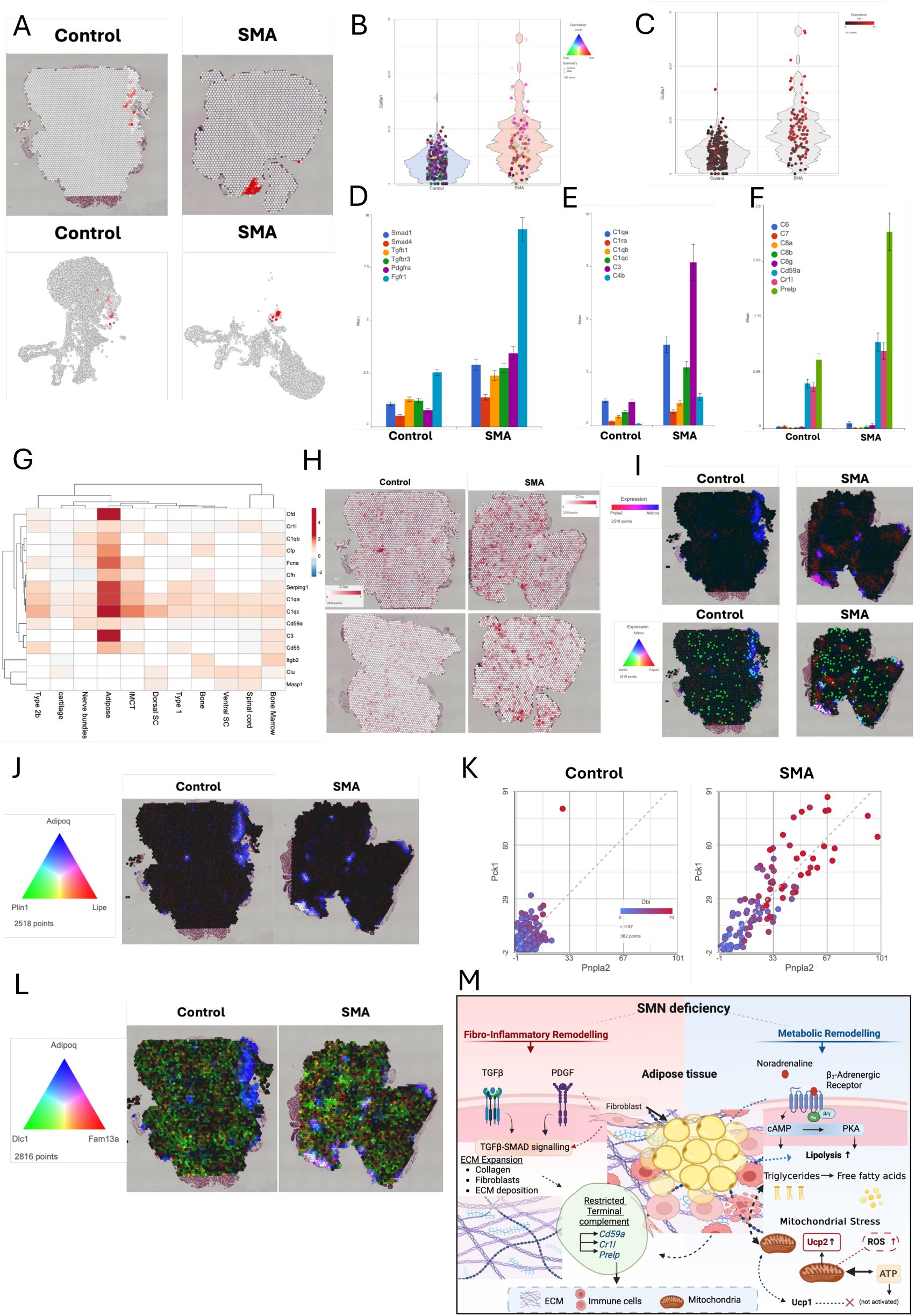
Fibro-inflammatory and metabolic remodeling in SMA adipose tissue. **A)** Spatial heatmap (top) and UMAP (bottom) showing UMI count expression for *Col6a1* in adipose tissue. Increasing red intensities indicate higher gene expression. **B)** Violin plots comparing *Col6a1* UMI gene expression (x-axis) between control and SMA adipose tissue. Expression of *Lama4* (blue), *Prelp* (Green), and *Nid1* (red) as a function of color intensity, where concurrent gene expression is white. **C)** Violin plot showing *Fgfr1* UMI gene expression of (red). Increasing red indicates higher expression. DESeq2 normalized mean gene expression of **D)** profibrotic signaling components (*Smad1, Smad4, Tgfb1, Tgfbr3, Pdgfra, Fgfr1*), indicating activation of TGFβ–SMAD and PDGF signaling pathways, and **E)** early complement pathway genes and **F)** terminal complement components (C5–C9) and complement regulatory genes (*Cd59a, Cr1l, Prelp*) in adipose (SEM). **G)** Hierarchical clustering gene expression heatmap of complement family members. Color scale represents fold-change expression in SMA over control tissue. **H)** Spatial heatmap showing UMI counts for complement gene *C1qc* across the spinal column tissue section. **I)** Spatial heatmap showing gene expression for adipose marker Adiponectin (*Adipoq*, blue), adipose triglyceride lipase (*Pnpla2*, red) and β3-adrenergic receptor (*Adrb3*, green). Increased color intensities correlate to increased gene expression and purple ST spots indicate co-expression of *Pnpla2* and *Adipoq*, while white ST spots indicate co-expression of *Pnpla2*, *Adrb3* and *Adipoq*. **J)** Spatial co-expression heatmap of adipocyte marker *Adipoq* (blue) and lipolysis regulators *Plin1* (green), *Lipe* (red). Increased color intensities correlate to increased gene expression. White ST spots indicate co-expression of *Plin1*, *Lipe* and *Adipoq*. **K)** Scatterplots showing positive correlations between *Pnpla2* (x-axis) and *Pck1* (y-axis, r ≈0.87) and *Dbi* (r ≈0.86) in adipose. Increased red intensities correlates to increased *Dbi* gene expression. **L)** Spatial co-expression heatmap of adipocyte marker *Adipoq* (blue) and RhoA-regulatory genes *Dlc1* (green), *Fam13a* (red) in control and SMA tissues. White ST spots indicate co-expression of *Dlc1*, *Fam13a* and *Adipoq*. **M)** Model illustrating how SMN deficiency drives coordinated fibro-inflammatory and metabolic remodeling in SMA white adipose tissue. Activation of TGFβ–SMAD and PDGF signaling promotes ECM expansion and early complement activation, while terminal complement signaling is restricted by *Cd59a*, *Cr1l*, and *Prelp*. In parallel, β3-adrenergic mediated cAMP-PKA signaling promotes lipolysis and mitochondrial stress (*Ucp2*) without activation of thermogenic (*Ucp1*) programs. Created in BioRender. Rietz, A. (2026) https://BioRender.com/3qmjfng

Early complement component genes *C1, C3,* and *C4b* were upregulated >2-fold in SMA adipose while terminal pathway genes *C5-C9* remained low and unchanged (Figure 5E,F). This was accompanied by selective induction of complement inhibitors including *Cd59a* (prevents membrane attack complex assembly), and *Cr1l* (suppresses C3/C5 convertase activity; Figure 5F). *Prelp* (∼3-fold upregulated), a complement-binding inhibitor, provides a mechanistic basis for restricted terminal complement activation despite robust upstream signaling.

SMA adipose exhibited a marked ∼5-fold increased expression of *Pnpla2* (ATGL, adipose triglyceride lipase), indicating elevated triglyceride hydrolysis. This was accompanied by upregulation of the β3-adrenergic receptor *Adrb3*, which triggers cAMP-mediated lipolytic signaling, suggesting heightened adrenergic stimulation in diseased adipose (Figure 5I). This is consistent with the increase in PKA/cAMP-regulated metabolic genes *Plin1, Lipe, Cpt1b,* and *Acadm* (Figure 5J) and mitochondrial cAMP targets *Ppargc1b* and *Prkar2b*. Upregulation of *Prkar2b* together with increased *Adrb3* and *Pnpla2* indicates enhanced cAMP-PKA signaling directed toward lipolysis rather than thermogenesis. Correlation analysis anchored to *Pnpla2* further revealed strong positive associations with *Pck1* and *Dbi* (r >0.86), which are involved in glyceroneogenesis and acyl-CoA handling, respectively (Figure 5K). This tight co-expression suggests coordinated activation of lipid mobilization and fatty-acid metabolic pathways in SMA adipocytes, consistent with stress-driven lipolysis. IPA metabolic and reactome pathway analysis of DEGs further supported this metabolic shift, identifying significant enrichment of pathways related to branched-chain amino acid degradation, fatty-acid β-oxidation, oxidative phosphorylation, and TCA cycle metabolism, consistent with enhanced mitochondrial substrate utilization (Figure S4D). IPA Reactome pathway analysis also highlighted activation of lipid mobilization and mitochondrial metabolic programs, including PPARα-regulated lipid metabolism, mitochondrial protein degradation, and Rho-GTPase signaling, indicating coordinated dysregulation of lipid handling, mitochondrial function, and stress-responsive signaling networks in SMA adipose (Figure S4E).

This signaling architecture aligns with increased *Ucp2* and absent *Ucp1* induction, indicating stress-driven metabolic activation rather than beige/thermogenic programming in SMA white adipose. *Ucp2* rises in response to lipolysis-induced mitochondrial stress, while persistent TGFβ–SMAD signaling suppresses adipocyte identity and prevents *Ucp1* activation. The upregulation of *Cyba* and *Arhgef1* reflects maladaptive activation of mechanosensitive and ROS-generating pathways that amplify RhoA-ROCK dependent cytoskeletal tension in SMA adipose, whereas increased *Dlc1* and *Fam13a* likely represent compensatory responses aimed at restraining excessive contractility due to lipolysis and mitigating metabolic and mitochondrial stress (Figure 5L,M; Table S11).

### Pathological alterations in the DRG

DESeq2 identified several genes with negative fold changes despite higher raw counts in SMA relative to control. Comparison of raw UMI counts (Figure S5A) and DESeq2 median-ratio normalized counts across spatial transcriptomic spots revealed discordance in relative expression between control and SMA DRG samples (Figure S5B). Normalized count distributions were compressed in ST spots with higher total RNA output, altering relative expression levels despite higher raw counts. Similar behavior was reported when the assumptions underlying median-ratio normalization - that most genes are not differentially expressed or that expression changes are balanced across conditions - are not met ^28,29^ ^30^. Counts that were log2-normalized showed distributions similar to raw UMI counts and did not exhibit fold-change inversion (Figures S5A,C). Subsequent differential expression analysis using the Hurdle model^31^ produced fold-change estimates that were directionally consistent with raw expression patterns and identified widespread gene upregulation in the DRG. This emphasizes the importance of validating normalized differential expression results against raw count distributions.

Hurdle analysis revealed coordinated upregulation of glutamate receptors, intracellular signaling pathways, and excitability-related genes, consistent with enhanced glutamatergic signaling (EGS) in SMA DRG neurons (Table S12). IPA indicated the EGS was the most significant pathway (Figure 6A). Both ionotropic and metabotropic glutamate receptors were represented, including AMPA receptor subunits *Gria1* and *Gria3*, the NMDA receptor subunit Grin3A, and group III metabotropic receptors Grm4 and Grm8, consistent with strengthened excitatory synaptic transmission and modulation of glutamate release (Figure 6B). Genes encoding calcium- and kinase-dependent signaling components downstream of glutamate receptor activation were also upregulated. Expression of the calcium-dependent kinase *Camk2a*, a key mediator of activity-dependent synaptic plasticity, increased by ∼3-fold, along with p38 MAPK pathway members *Mapk11* (p38β) and *Mapk12* (p38γ), which are implicated in pain sensitization^32^ (Figure 6B). The non-receptor tyrosine kinase *Src*, known to potentiate NMDA and AMPA receptor function through phosphorylation, further supports enhanced glutamatergic signaling. Multiple phospholipase and second messenger signaling genes were differentially expressed including *Plcb1*, *Plcb4*, and *Plcg1*, consistent with enhanced coupling of metabotropic glutamate receptors to IP_3_/DAG-mediated calcium signaling. In parallel, lipid metabolism genes *Pla2g12a*, *Pla2g4e*, *Pld2*, *Pld5*, *Abhd3*, *Napepld*, *and Pnpla2* suggest engagement of lipid-based neuromodulatory pathways that interact with glutamate signaling to regulate synaptic strength and neurotransmitter release (Figure 6B). Differential expression of *Atf2* and *Creb5* indicates activation of activity-dependent transcriptional programs associated with neuronal plasticity. Consistent with increased excitatory drive, voltage-gated sodium channel genes *Scn2a*, *Scn8a*, and the auxiliary subunit *Scn2b* were increased, supporting enhanced neuronal excitability. Changes in *Shank2*, a postsynaptic glutamate receptor scaffold, and *Slc38a1*, a glutamine transporter supporting glutamate synthesis, suggest structural and metabolic adaptations may reinforce sustained glutamatergic transmission in SMA DRG. Collectively, dysregulation of glutamate receptors, downstream calcium- and lipid-dependent signaling pathways, and excitability-associated ion channels in SMA DRG neurons provides a plausible mechanistic basis for the heightened pain sensitivity observed in SMA patients and animals^33–35^. Additional pathways significantly dysregulated in SMA DRGs were retinoic acid receptors (RAR) activation, docosahexaenoic acid (DHA) signaling and synaptogenesis signaling. IPA Reactome analysis identified chromatin modifying enzymes as most significantly altered (Figure S6). Consistent with this enrichment, key chromatin and transcriptional regulators including activity-dependent transcription factor *Atf2*, histone methyltransferases *Kmt2a, Kmt2b, Setd2, and Dot1l*, and histone acetylation and co-activation components K*at6b, Brd8, Tada2b, and Ncoa1*, as well as regulatory factors such as *Hdac10* and Jak2 were differentially expressed, which indicates epigenetic reprogramming represents a prominent transcriptional signature in SMA dorsal root ganglia.

**Figure 6.**
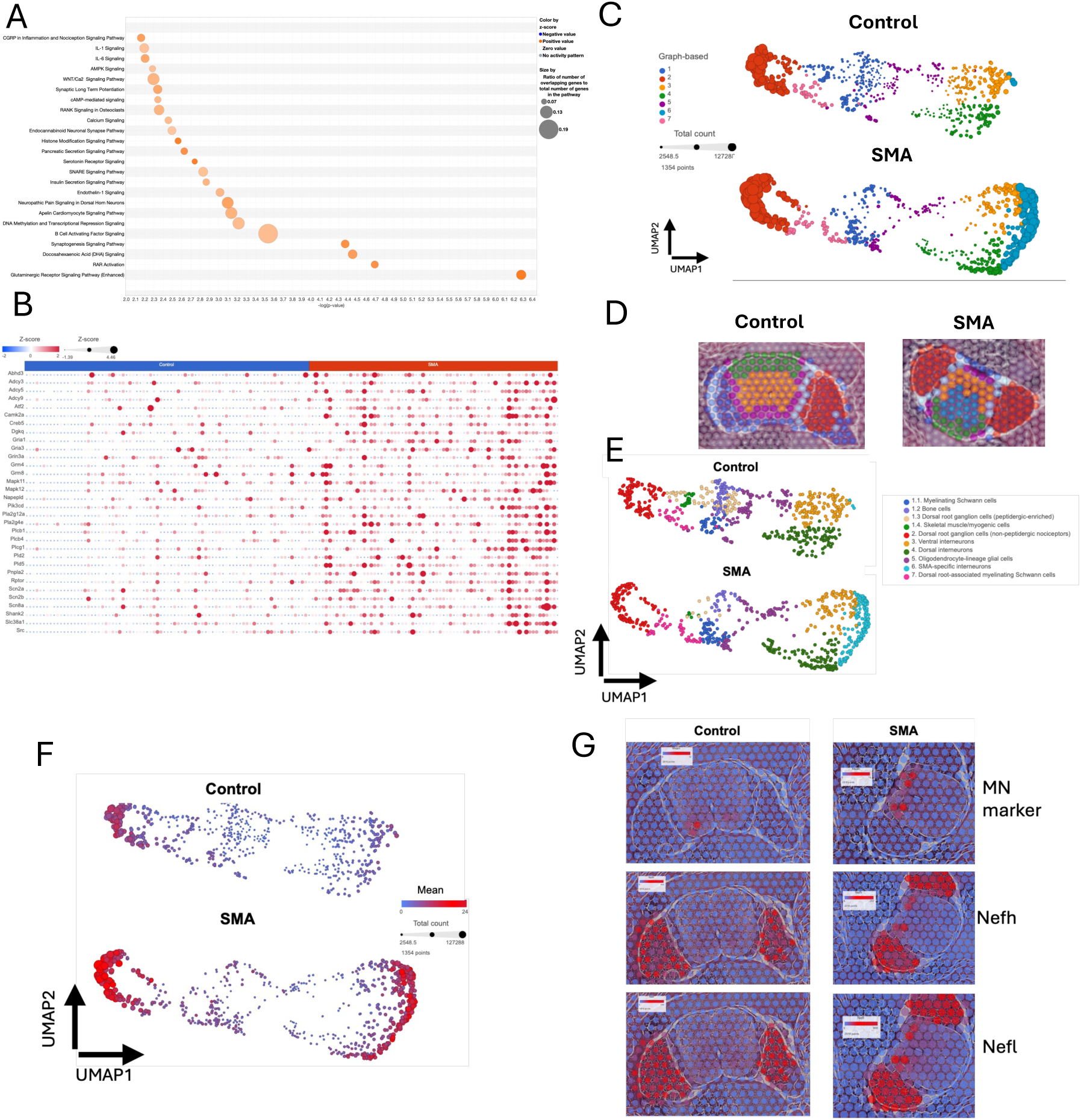
SMA drives spatially patterned transcriptional reprogramming across the spinal cord. **A)** IPA of DEGs identified in SMA DRG highlighting the top 20 enriched signaling pathways. **B)** Heatmap analysis of raw UMI counts in DRG ST spots for DEGs identified in the glutamatergic signaling pathway. **C)** UMAP of ST spots from spinal cord sections, colored by graph-based clustering and size adjusted for total UMI count, demonstrating a significant increase in transcript count in SMA specific cluster 6. **D)** Insets show the corresponding spatial localization of clusters overlaid on tissue sections, highlighting preservation of major anatomical domains. **E)** UMAP of ST spots annotated by inferred cell-type identities, including DRG neurons, interneurons, Schwann cells, and an SMA-specific cluster present exclusively in SMA samples. **(F)** UMAP of spinal cord with heatmap showing mean expression of RNA-processing genes (*Srsf11, Sf1, Snrpa1, Clasrp,* and *Rsrp1*) across dorsal, ventral, and DRG neuronal populations. Red indicates higher and blue lower expression, highlighting coordinated spliceosomal upregulation in SMA neurons and DRG. **(G)** Spatial heatmap depicting increased expression of neurofilament genes (*Nefh* and *Nefl*) in ventral SMA spinal cord. Elevated expression localizes to regions enriched for cholinergic motor neuron markers *Chat, Slc18a3, Mnx1,* and *Uts2* in SMA tissue. Red indicates higher and blue lower expression.

### Pathological alterations in the spinal cord highlight effects on extracellular matrix (ECM)

While primarily considered a disease of alpha motor neurons, experimental data implicate sensory neurons, interneurons, astrocytes, and muscle in pathogenesis of SMA. The sub-clustering analysis of the spinal cord resolved multiple clusters defined by their anatomical region (Figure 6C). Of particular interest was the emergence of a unique SMA-specific cluster (Cluster 6C,D) localized within the lateral and extending into ventral horn where motor neurons reside. Cluster 6 exhibited increased transcriptional activity, characterized by both an increased number of total transcripts per spot and an increased number of expressed genes (Figure 6D, Figure S7A). Cluster 6 expressed inhibitory and excitatory neuron markers *Slc32a1, Slc6a7 and Slc17a6* together with developmental and caudal identity genes *Neurod6, Unc5a, Hoxc11, and Otp*, consistent with the emergence of a disease-associated lateral/ventral interneuron population.

Cluster 1 was initially identified as a heterogeneous non-neuronal compartment adjacent to the dorsal root ganglion and spinal column, enriched for extracellular matrix and collagen-associated genes. Subclustering of Cluster 1 resolved four distinct transcriptional populations corresponding to discrete anatomical compartments within the spatial field. One subcluster expressed canonical myelinating Schwann cell markers *Mbp, Mag, Plp, Ugt8a, and Fa2h* together with myelination-associated transcription factors *Pou3f1 and Egr2*, indicative of peripheral nerve associated Schwann cells. A second subcluster was defined by strong expression of osteogenic and bone matrix genes *Ibsp, Bglap, Dmp1, Sp7, and Phex* and collagen-rich extracellular matrix components *Col1a1, Col1a2, Pcolce, and Serpinh1*, identifying periosteal or bone-associated osteogenic lineage cells. A third subcluster expressed DRG sensory neuron markers *Pirt, Scn10a, Prph, and Calca*. The fourth subcluster showed robust expression of skeletal muscle and myogenic genes *Myf6, Myh3, Casq2, Ldb3, and Smpx*, consistent with adjacent muscle tissue (Figure 6E). These findings demonstrate that Cluster 1 represents a composite anatomical region comprising peripheral nerve, bone cells, sensory neurons, and skeletal muscle.

Cluster 2 is enriched for canonical markers of non-peptidergic nociceptors including *Mrgprd/Mrgprx1, P2rx3, Scn11a,* and *Isl2*, suggesting that this population is predominantly composed of non-peptidergic DRG sensory neurons. While both cluster 1.3 and cluster 2 represent DRG sensory neurons, cluster 1.3 is enriched for peptidergic nociceptor markers *Calca, Scn10a and Pirt*, whereas cluster 2 predominantly expresses non-peptidergic nociceptor markers *Mrgprd/Mrgprx1, P2rx3, and Scn11a*, indicating capture of distinct sensory neuron subtypes with differing spatial distributions. Cluster 7 represents a distinct transcriptional cluster adjacent to the DRG that expresses canonical myelinating Schwann cell markers *Prx, Pmp2, Mal, Cldn19, Gjc3, and Cdh19*, consistent with peripheral nerve tissue of the dorsal root. While both clusters 1.1 and 7 represent myelinating Schwann cells, cluster 1.1 displays a canonical myelin gene program characteristic of peripheral nerve Schwann cells, and cluster 7 is enriched for sensory neuron associated transcripts *Pirt, Scn10a, Scn11a, and Calca*, suggesting a population of dorsal root associated myelinating Schwann cells with enhanced axon glia coupling. Cluster 1.3 (peptidergic DRG neurons) was represented by substantially fewer ST spots in SMA compared to control tissue and was excluded from downstream analyses.

Based on expression of canonical glycinergic and caudal spinal markers (*Slc6a5/Glyt2, Slc7a10, Hoxc12, Hoxc13, Cbln4)* together with synaptic and membrane-associated genes (*Tmem91, Olfm3, Cst3, Elmod1, Ubap1l)*, Cluster 3 corresponds to ventrally located interneurons. Cluster 4 expressed dorsal spinal interneuron markers *Lmx1b, Lbx1, and Ebf2*, consistent with its anatomical location. Cluster 5 expressed canonical oligodendrocyte and CNS myelin markers *Plp1, Cnp, and Qk* together with additional glial-associated genes *Cd81, Atp1a2, and Aqp4*. Collectively, these clusters recapitulate the known dorsoventral organization of spinal cord cell types.

Trajectory analysis (Monocle 3^36^) of spatial transcriptomic spots rooted in ventral and dorsal spinal domains revealed a disease-associated extension of pseudotime in SMA, with control spots confined to a restricted range while the SMA-specific cluster extended into late trajectory states (Figure S7B-J). This cluster was absent from control tissue and localized almost entirely to maximal pseudotime (95% of spots), with 65% of all SMA ventral spots occupying this terminal transcriptional state. Genes positively correlated with pseudotime, including positional identity genes *Hoxc8, and Hoxc*9, neuronal differentiation factor Neurod6, chromatin regulators *Hdac9, Phf13, Taz, and Dcaf15*, synaptic genes *Kcnh7, Mast3, Pcdha12, and Cpne7*, and protein homeostasis components *Atg16l2, Edem2, and Usp29*, were enriched in this cluster, indicating that SMA drives neurons toward a late pathological state characterized by transcriptional reprogramming, synaptic remodeling, and dysregulated regional identity (Figures S8A,B). Given the extension of the SMA specific cluster outwards from both dorsal and ventral interneuron populations in the UMAP plots, we performed subclustering to resolve its cellular composition. This analysis identified dorsally and ventrally derived neuronal subpopulations within the SMA specific cluster (Figures S9A, B). Biomarker profiling confirmed expression of canonical excitatory and inhibitory markers across control and SMA dorsal and ventral spatial transcriptomic populations (Figure S9A). DEGs were stratified into region-specific and shared categories, which identified 172 ventral-specific DEGs and 475 dorsal-specific DEGs, compared with 110 DEGs common to both anatomical compartments (Tables S13, S14). Shared dorsal and ventral gene expression changes included coordinated upregulation of multiple components of the RNA-processing and spliceosomal machinery, including *Srsf11, Sf1, Snrpa1, Clasrp, and Rsrp1* (Figure 6F). Given the role of SMN in snRNP assembly and spliceosome function, enrichment of these core splicing regulators is consistent with altered spliceosomal homeostasis in SMN-deficient neurons. UMAP visualization further revealed significant upregulation of these splicing factors within DRG neuronal populations (Figure 6F). IPA of dorsal SMA interneurons identified multiple dysregulated pathways associated with synapse biology (Figure S9C). Core presynaptic components including *Syn2* and *Syt7* together with synaptic scaffold and signaling regulators such as *Shank3, Kalrn, and Git1*, were significantly elevated. Multiple voltage-gated calcium channel subunits were upregulated, including *Cacna1h, Cacna1c, Cacna1d*, *Cacna1e*, *Cacna1a*, and the auxiliary subunit *Cacna2d1*. In parallel, several members of the calcium-dependent membrane-binding copine family (*Cpne1, Cpne2, Cpne7,* and *Cpne8*) were coordinately increased, supporting augmented calcium-responsive membrane dynamics and vesicular trafficking. These transcriptional changes were confined to regions enriched for canonical cholinergic motor neuron markers including *Chat, Slc18a3, Mnx1*, and *Uts2,* consistent with the histology (Figure 6G). Although fewer DEGs were identified in the ventral SMA interneuron cluster compared to the dorsal compartment, the ventral signature was distinguished by prominent dysregulation of cytoskeletal genes. Neurofilament family members *Nefm*, *Nefh*, and *Nefl* exhibited two- to three-fold increases, whereas *Ina* expression remained unchanged. These neurofilament genes did not show comparable upregulation in the dorsal compartment (Figure 6G, Figure S9D).

IPA of ventral SMA DEGs identified enrichment of pathways involved in cytoskeletal structure, vesicle trafficking, and calcium-dependent synaptic regulation (Figure S9E). The RHO GTPase cycle emerged as a central node, supported by upregulation of actin regulators *Arhgef28, Farp1, and Rhou* and cytoskeletal effectors (e.g. *Stmn2*). Concurrent enrichment of microtubule-based transport pathways aligned with increased expression of *Kif3a, Dync1h1*, and *Dctn1* together with induction of neurofilament genes *Nefm, Nefh, and Nefl*. Vesicle-associated genes *Syn2 and Syt7* and voltage-gated calcium channel subunits further support activity-dependent changes.

Angiotensinogen (*Agt*) expression increased approximately four-fold in ventral SMA neurons (Figure S9F), suggesting engagement of local renin-angiotensin system (RAS) signaling in the presymptomatic spinal cord. Consistent with this, multiple downstream RAS-associated signaling components, including *Adcy3, Plcg1,* and *Mapk* pathway members, were concurrently elevated in several tissues (Figure S9G). Central RAS activation has been implicated in neuronal stress responses, modulation of calcium signaling, and oxidative stress pathways, and is also observed following spinal cord injury, where it contributes to early injury associated signaling cascades. Although overt inflammatory signatures were not detected, increased *Agt* expression together with coordinated MAPK/PI3K pathway engagement may reflect a subtle injury like stress program within ventral motor-associated regions.

SMA neurons exhibited coordinated upregulation of several tubulin isoforms (*Tuba1a, Tubb2a, Tubb2b, Tubb3, Tubgcp6*) in both ventral and dorsal spinal cord compartments (Figure S9G). Given the central role of microtubules in axonal integrity and intracellular transport, these changes are consistent with cytoskeletal instability and an early axonal stress response in presymptomatic SMA. In parallel, major fibrillar collagens *Col2a1, Col1a1, and Col1a2* were downregulated in the ventral region (Figure 7A, B), indicating selective extracellular matrix reorganization. Correlation analysis anchored to Col1a1 expression in the ventral compartment revealed limited coordination within the extracellular matrix program. While *Col1a1* and *Col1a2* were strongly co-expressed (ρ ≈ 0.81), Col3a1 (ρ ≈ 0.56) and Col5a2 (ρ ≈ 0.52) demonstrated moderate positive associations. Other fibrillar collagens and key matrix regulators such as *Lox, Serpinh1, Pcolce*, and *Mmp2* showed weak correlations (ρ < 0.5), suggesting that collagen deposition is not tightly coupled to crosslinking, chaperone activity, or proteolytic alterations at the transcriptional level within ventral tissue. In contrast, the tubulin-anchored analysis revealed a highly coherent intracellular remodeling program. *Tuba1a* exhibited extremely strong correlations with additional tubulin isoforms *Tubb2a, Tubb2b,* and *Tubb3*, microtubule regulators *Stmn3* and *Dpysl3*), membrane–cytoskeletal adaptor *Sptbn1*, and axonal growth-associated genes *Gap43* and *Marcksl1*). *Stmn3*, a direct regulator of tubulin polymerization dynamics, showed one of the strongest associations with *Tuba1a* (ρ ≈ 0.93; Figure 7C). Unlike the ECM program, this cytoskeletal module displayed uniformly high correlations across both structural components and regulatory effectors. Fold-change heatmaps of collagen and other ECM proteins across tissues reveal heterogeneous, tissue-dependent regulation of collagen and ECM genes in SMA (Figure 7D, Figure S10, Table S16, S17). Rather than a uniform shift, distinct collagen isoforms and matrix components are selectively up- or down-regulated across spinal cord, adipose, muscle, and other tissues, indicating context-dependent changes of the extracellular scaffold. Concomitant dysregulation of ankyrin family members (Figure 7E, Table S18), which are key adaptors linking adhesion molecules such as *L1cam* to the spectrin-actin cytoskeleton, suggests that extracellular changes are coupled to altered membrane-cytoskeletal anchoring across tissues (Figure 7C). Together, these analyses indicate that ventral SMA spinal cord tissue engages an integrated intracellular cytoskeletal remodeling program, whereas extracellular matrix changes appear more heterogeneous and potentially governed by additional spatial or post-transcriptional mechanisms.

**Figure 7.**
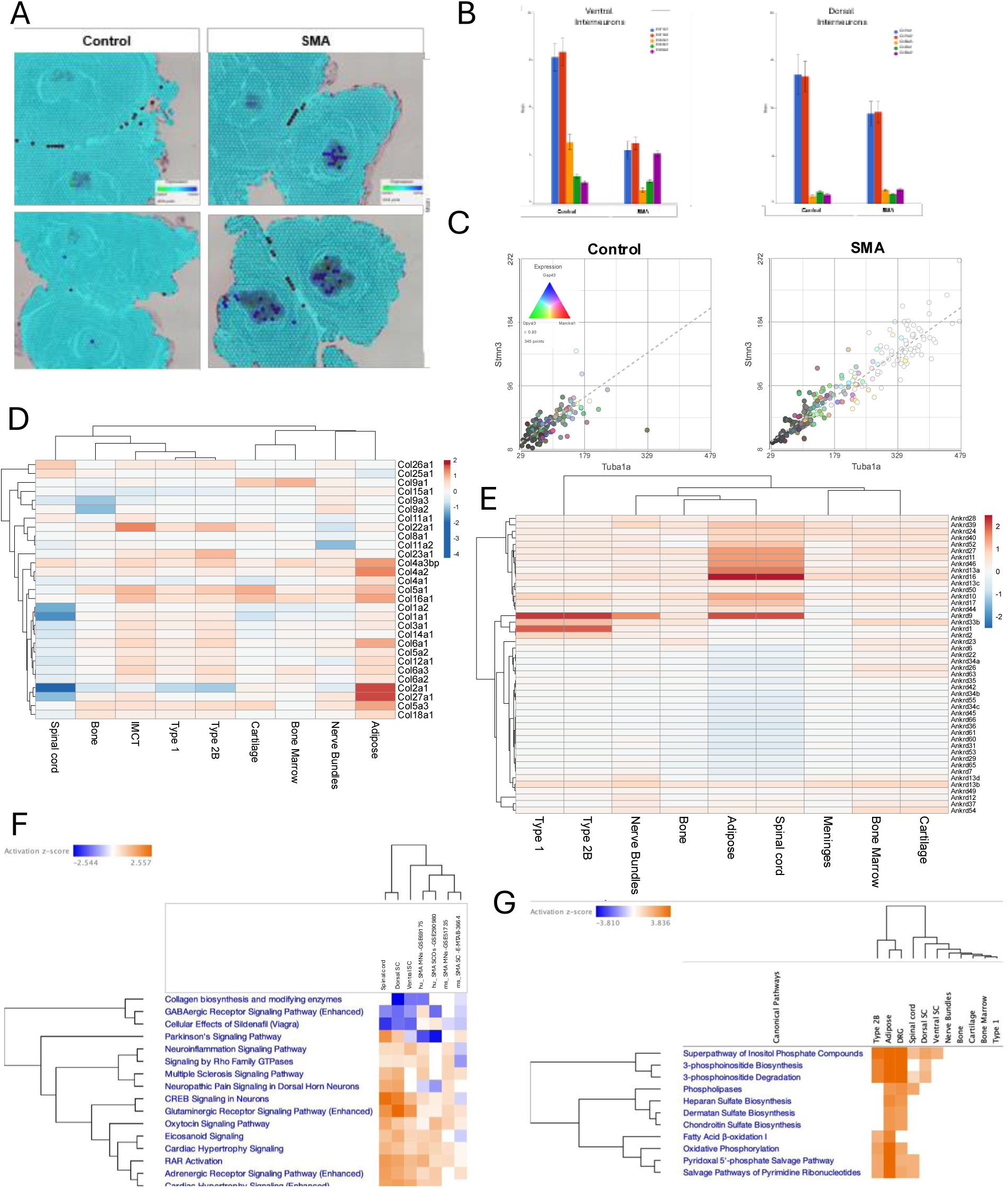
Pathway alterations in SMA spinal cord and peripheral tissues. **(A)** Spatial heatmap analysis of raw UMI counts for *Col1a1* (green) and *Col1a2* (blue) in spinal column sections. Cyan indicates co-expression. SMA tissue shows lower expression of *Col1a1* and *Col1a2* in ventral regions compared to control. **B)** Bar plots show relative proportions of DESeq2 normalized mean counts for several collagen genes in ventral interneurons (left) in comparison to their expression the dorsal SMA cluster (right). **C)** Correlation of *Tuba1a* with *Stmn3* expression in the ventral compartment (ρ ≈ 0.93). Coloring by *Gap43* (blue), *Dpysl3* (green), and *Marcksl1* (red) highlights joint enrichment (white dots) of axonal growth and cytoskeletal regulators along the high *Tuba1a/Stmn3* axis, consistent with coupled microtubule and neurite remodeling in presymptomatic SMA. Hierarchical clustering heatmap of **D)** collagen family members and **E)** ankyrin family members across annotated tissue regions. Color scale represents fold-change expression in SMA tissue. **F)** Activation analysis (z-score) of select signaling pathways comparing RNA expression from our ST data set to RNA seq data from published murine PND5 SMA spinal cord (E-MTAB-3664), human (GSE69175), murine (GSE51735) SMA motor neurons, and human spinal cord organoid (SCO; GSE290980). **G)** Metabolic and biosynthetic pathway activation analysis (z-score ≥ 2) across SMA tissue regions from this study.

Comparative analysis demonstrated strong concordance between our spatial transcriptomic data and published RNA-seq results including activation of neuronal signaling pathways (CREB, glutamatergic, Rho GTPase, and adrenergic signaling) and dysregulation of collagen pathways (Figure 7F). Metabolic pathway analysis revealed tissue-specific but convergent activation of lipid and phosphoinositide signaling across several SMA tissues (Figure 7G).

## DISCUSSION

Application of spatial transcriptomics (ST) to cross-sections from the entire lumbar region of pre-symptomatic SMA mice defined early, tissue-wide transcriptional changes by preserving spatial context and capturing cells that may be underrepresented with dissociative approaches. ST allowed characterization of gene expression in muscle, bone, cartilage, adipose tissue, and bone marrow.

Our ST analysis reveals that SMA skeletal muscle undergoes early, fiber type-specific transcriptional alterations characterized by activation of stress-responsive and atrophy-associated gene programs prior to reported overt degeneration in this SMA mouse model^37^. Both Type 1 and Type 2b muscle fibers exhibit induction of canonical atrophy-related genes including *Trim63, Fbxo32, Fbxo40*, and *Foxo* family members, alongside stress and metabolic regulators, indicating engagement of catabolic pathways. These changes occurred earlier than typically reported in SMA models, whereas muscle atrophy and activation of ubiquitin–proteasome and autophagy pathways are generally associated with later stages of disease progression and denervation. This reveals that activation of atrophy-associated transcriptional programs represents an intrinsic and early event rather than a secondary consequence of motor neuron loss. The concurrent upregulation of oxidative, mitochondrial, and hypertrophic remodeling pathways (e.g., *Nmrk2, Perm1, Abra, Mph*) indicates the presence of compensatory mechanisms aimed at maintaining energy homeostasis and contractile function. The stronger transcriptional perturbation observed in Type 2b fibers is consistent with their known vulnerability in neuromuscular disorders^38^. In addition, alterations of neddylation and RNA-processing pathways including direct SMN-interacting proteins, highlights potential mechanistic links between SMN deficiency and muscle dysfunction.

Bone in SMA is fragile and prone to fracturing ^39–41^, especially in later-onset forms of SMA, and is attributed to reduced muscle function and impaired weight-bearing leading to low bone mineral density^26,42,43^. Abnormal skeletal development can result in scoliosis, hip instability, and long-bone deformities, with scoliosis being a common complication^19,21,24^. Scoliosis is being detected in a subset of treated infants despite maintenance of motor function^19–26^. While some recent studies report intrinsic bone changes in SMA, to our knowledge our ST analysis is the first in calcified neonatal murine bone and reveals early molecular dysregulation at a pre-symptomatic stage. We identified a protease-rich, osteoclast-oriented transcriptional program characterized by increased expression of *Acp5, Ctsk, Mmp9, and Mmp13*, and osteopontin (*Spp1*), consistent with enhanced bone resorption and osteoporosis^44, 45,46^. This aligns with reports of elevated serum osteopontin levels in SMA patients^47^ and late-stage SMA mice^48^, increased cathepsin activity in neurodegeneration^49^, which together may contribute to fracture risk in SMA. The detection of increased osteoclastogenic and protease programs at PND4 suggests that skeletal dysregulation precedes motor neuron deficits in SMA^50^.

Spatial profiling revealed an active bone marrow program characterized by granulopoiesis, degranulation, and acute-phase signaling, forming a feed-forward niche that could promote osteoclast activity and matrix loss. Although induction of regulatory genes (e.g. *Zfp36l2, Cst3, Fst*) suggests compensatory restraint, the balance remains catabolic at this pre-symptomatic stage. Together, these findings suggest that intrinsic skeletal alterations, including potential cartilage changes such as reduced *Acan* expression, may contribute to skeletal fragility independent of muscle weakness.

Individuals with SMA exhibit characteristic alterations in body composition, including increased fat mass relative to lean mass, even at low body weight, along with evidence of systemic metabolic dysfunction such as dyslipidemia and impaired glucose tolerance. Our data indicate that SMA engages pathways driving adipose and metabolic dysregulation^51–53^. Indeed, we observed that adipose alterations in presymptomatic SMA animals reflects stress-induced lipolysis driven by metabolic and inflammatory signaling. Upregulation of *Pnpla8* indicates mitochondrial strain associated with excess fatty-acid flux. Concurrent ECM dysregulation, complement activation, and pro-fibrotic signaling, together with adrenergic lipolysis, may disrupt adipocyte identity and promote oxidative stress, consistent with increased mitochondrial markers such as *Atp5a1*. These findings define a dysfunctional adipose microenvironment characterized by inflammation and metabolic imbalance. These alterations mirror broader systemic features of SMA including impaired lipid handling and chronic inflammation and are supported by a 12-fold increase in Haptoglobin (*Hp*) gene expression. Although the macrophage marker *Adgre1* was unchanged, immune infiltration may be underestimated due to the spatial resolution of the Visium platform. Pathway analysis revealed widespread metabolic dysregulation, including TCA cycle, triglyceride, inositol, and amino acid metabolism (Figure S4D), along with enrichment of Rho GTPase signaling. Genes associated with ECM remodeling and inflammation (*Arhgef1, Rock2, Net1*, Table S11) were upregulated alongside compensatory regulators (*Fam13a, Dlc1*, Figure S4F), consistent with reported abnormalities in lipid metabolism and mitochondrial function^54^.

Consistent with broader immune dysregulation in SMA, early complement activation in adipose tissue likely promotes chronic stromal inflammation. Complement components C1q and C3 are upregulated across multiple tissues including muscle, spinal cord, bone marrow, and DRG, indicating systemic yet tissue-specific complement involvement. SMN deficiency disrupts both innate and adaptive immunity, with prior studies reporting altered cytokine profiles, impaired immune cell function, and increased inflammation^55;54;56–58^, which may exacerbate neuronal vulnerability^59^. Our data further show selective activation of early complement pathways in adipose tissue with restriction of terminal complement-mediated cytolysis, favoring chronic inflammatory and fibrotic signaling. Supporting this, *Prelp*, which was upregulated in SMA adipose, has been linked to fibrosis and inflammation in metabolic disease models^60^. Complement activation has been implicated in SMA CNS pathology, where C1q and C3 drive microglia-dependent synapse loss^61^. In addition to adipose tissue, *C1qa, C1qb*, and *C1qc* mRNAs were significantly increased in multiple SMA tissues, including muscle, spinal cord, bone marrow, and DRG, with weaker induction in bone and nerve bundles, while cartilage showed no significant changes. Similarly, C3 induction was restricted to specific tissues, underscoring the tissue-dependent regulation of complement signaling in SMA (Figure 5G,H; Table S15).

Our findings extend SMA beyond motor neuron degeneration to reveal coordinated, multi-cellular dysregulation across the spinal cord and peripheral compartments. ST identified a disease-specific neuronal population of dorsally and ventrally derived interneurons that occupies a terminal transcriptional state and localizes to lateral and ventral horn regions. This population is enriched for positional identity, chromatin, and synaptic pathways, consistent with convergence toward a late pathological state. SMA spinal cord regions were characterized by both shared and region-specific responses, including widespread dysregulation of RNA processing and spliceosomal machinery, consistent with the role of SMN in snRNP assembly^62^. Dorsal regions were enriched for synaptic and excitability-related programs, whereas ventral regions showed prominent cytoskeletal alterations, with neurofilament upregulation restricted to motor neuron regions and broader tubulin dysregulation across ventral tissue. These changes were accompanied by activation of Rho GTPase and microtubule transport pathways, consistent with early axonal stress and structural adaptation. Dorsal and ventral compartments showed differences in pathway magnitude but preserved a core SMA transcriptional signature consistent with prior RNA-seq studies. These data suggest that SMN deficiency drives early, spatially organized transcriptional reprogramming linking RNA processing, synaptic dysfunction, and cytoskeletal remodeling to a disease-specific neuronal state prior to overt neurodegeneration. Enrichment of chromatin regulators in the SMA-specific state highlights epigenetic modulation as a potential therapeutic strategy, while osteoclastogenic and marrow inflammatory programs suggest targets for early skeletal pathology. The spatially localized disease signatures identified here provide a framework for targeted validation of therapeutic intervention.

## Conclusions

Spatial transcriptomics of pre-symptomatic SMA mice reveals that SMN deficiency triggers a coordinated, multi-tissue pathological program encompassing neuronal transcriptional reprogramming, cytoskeletal alteration, synaptic and excitability adaptations, along with marrow inflammation, and tissue-specific ECM and metabolic changes. These linked processes form spatially defined microenvironments that likely contribute to early dysfunction and drive downstream degeneration. Interventions that combine SMN restoration with approaches that stabilize cytoskeletal and synaptic integrity, temper maladaptive activity, and mitigate marrow, skeletal and metabolic stress may therefore have the greatest potential to alter disease trajectory.

## Limitations

The mouse model of SMA may not accurately reflect human disease, however pre-symptomatic and pre-morbid spinal cord from humans cannot be similarly investigated. Low levels of SMN are reported to induce alterations in mRNA splicing, which are not detectable with the short sequences produced in ST. Heart, liver, spleen, CNS including cerebellum, were not sampled and mRNA expression may not accurately parallel protein levels and activity.

## Acknowledgements

This project was supported by grants from the Ralph W. and Grace M. Showalter Research Trust (AR) and NIH R01NS082284 (EJA) and the Indiana Clinical and Translational Sciences Institute (UL1TR002529). The content is solely the responsibility of the authors and does not represent the official views of the National Institutes of Health or Indiana University.

To support the interpretation of results, assist in drafting sections of the manuscript, and help identify potential errors or inconsistencies, we made use of ChatGPT5. All final interpretations, decisions and conclusions remain our own and we take full responsibility for the content of this publication.

## METHODS

### Mice breeding, genotyping, and treatments

The animal study protocol was approved by the Institutional Animal Care and Use Committee of Indiana University and conformed to the Guide for the Care and Use of Laboratory Animals. We used the congenic severe ‘Li’ (Taiwanese) SMA mouse model (Jax #5058; FVB.Cg-*Smn1*^tm1Hung^Tg(SMN2)2Hung/J)^63^. Their breeding scheme results in 50% of the litter developing the SMA-like phenotype and 50% healthy heterozygous (Het) siblings. Severe SMA (5058) neonatal mice were bred as previously reported^64^. Animals were maintained on a 12 to 12 h light/dark cycle with food and water ad libitum and were provided with Bed-r’Nest as standard of care. Neonatal mice were cryo-anesthetized ^65^ and sacrificed on PND4. L4–L5 spinal columns were extracted for subsequent formalin fixation, while in parallel DNA was isolated from tail snips (0.1–0.15 cm) using the QIAGEN DNeasy Kit. SMN1^tm^^1^ and SMN2 and Y Chromosome (*Sry*) genotyping was performed according to the protocol provided by Jackson Laboratory.

### Preparation of tissue sections for Visium

Formalin-fixed L4-L5 spinal columns were paraffin-embedded (FFPE) according to genotype and sex. Spinal columns from two animals of the same genotype and sex were embedded together. Here we used only female control and SMA animals. Tissue blocks were sectioned at 5 µm thickness at the Indiana University (IU) Histology Core. Prior to preparation of the Visium slides, we validated the integrity of the FFPE tissue sections via hematoxylin and eosin (H&E) staining and extracted RNA from adjacent tissue sections to measure the RNA quality with the Agilent Bio-analyzer 2100 and the Eukaryote Total RNA Pico assay v2.6 ^66^. All RNA samples had a DV200 ≥ 90% (Figure S1A, Table S1), which far exceeded the recommended DV200 ≥ 50%.

### RNA isolation for cDNA library generation and sequencing

Tissue sections were placed onto Visium slides using the 10x Genomics CytAssist system. Spatial transcriptomics library preparation, sequencing, and primary data processing were performed at the Indiana University School of Medicine Medical Genomics Core and the Center for Computational Biology and Bioinformatics according to the manufacturer’s protocol (Visium Spatial Gene Expression for FFPE, 10x Genomics).

Sequencing was performed on an Illumina NovaSeq 6000 platform (Center for Medical Genomics, Indiana University School of Medicine) to generate 28-bp reads containing spatial barcode and UMI sequences and 50-bp probe reads. Raw sequencing data were processed using Space Ranger v2.1.1 (10x Genomics), incorporating a modified mouse probe set (Visium_Mouse_Transcriptome_Probe_Set_v1.0_mm10-2020-A.csv) to enable detection of Pls3 (Plastin 3). Briefly, spaceranger mkfastq (which wraps Illumina bcl2fastq) was used to demultiplex binary base call (BCL) files into sample-specific FASTQ files. Brightfield H&E and CytAssist images were manually aligned using the 10x Genomics Loupe Browser.

FASTQ files, aligned image data, and the corresponding JSON alignment files were processed using spaceranger count for sequence alignment, tissue detection, fiducial alignment, and spatial barcode/UMI quantification. Reads were aligned to the mouse reference genome (mm10) using the modified 10x Genomics whole-transcriptome probe reference for FFPE samples. Gene expression levels were quantified as unique molecular identifier (UMI) counts per spatial spot. Quality control metrics generated by Space Ranger are provided in Table S1.

### Spatial transcriptomic data processing and analysis

Partek® Flow was used for downstream normalization, visualization, and statistical analyses. Space Ranger outputs, including graph-based clustering and UMAP coordinates, were imported into Partek® Flow (Build version v12.9.1) for further analysis. Prior to analysis, boundaries between tissue sections were manually defined in Partek® Flow and annotated by individual animal ID (Control vs. SMA: D2, D4, E2, E8 vs. D3, D9, E9, E11).

Low-quality spots, including those located outside of tissue regions with low UMI counts or with less than 500 detected genes were excluded during quality control. To account for mixed cell signals inherent to ST spots, tissue clusters were manually annotated based on canonical marker gene expression, allowing assignment of spots to tissue types and minimizing the impact of overlapping anatomical regions. The number of detected genes and unique molecular identifiers (UMIs) per spot were assessed for each tissue section (Figures S1B, C).

Read counts were normalized using counts per million (CPM) and log-transformed as log2 (CPM +1). The log2 normalized data were used for principal component analysis (PCA), UMAP visualization, graph-based clustering, pseudotime trajectory analysis and biomarker gene identification. Highly variable features (n = 2,000) were identified for dimensionality reduction. PCA was performed, retaining 100 principal components, and graph-based clustering with biomarker analysis was conducted using the Louvain algorithm based on the top 10 principal components. For differential expression analysis, raw counts were used with DESeq2 normalization.

### Differential gene expression analysis

Differential gene expression (DEG) analysis was performed using DESeq2. To avoid zero counts, a value of 0.001 was added to all raw gene expression counts prior to analysis. Size factor normalization was conducted using the median-of-ratios method, followed by differential expression testing with DESeq2. Genes with low expression (mean expression ≤ 0.5) were excluded from the analysis. DEGs were defined as those with a false discovery rate (FDR) < 0.05 and an absolute fold change greater than 2.

DEG analysis of dorsal root ganglia (DRG) spatial transcriptomic data was initially performed using DESeq2; however, inconsistencies between raw UMI counts and normalized fold changes were observed, likely due to violation of median ratio normalization assumptions in samples with variable RNA content. To address this, DEG analysis was subsequently performed using the Hurdle model, which accounts for zero inflation and separates detection from expression magnitude. This approach produced fold-change estimates consistent with raw expression patterns and was used for downstream interpretation. Using the same thresholds as DESeq2, DEGs were defined as those with a false discovery rate (FDR) < 0.05 and an absolute fold change greater than 2.

### Pathway and downstream analysis

Ingenuity Pathway Analysis (IPA; QIAGEN) was performed using DESeq2 differential expression results, except for DRG cluster, where the Hurdle results were used. Genes with a false discovery rate (FDR) < 0.05 and an absolute fold change ≥2 were included. Canonical pathways with an absolute activation z-score ≥2 were considered significantly enriched. IPA was also used to perform comparative pathway analysis to the following published SMA RNA sequencing from murine PND5 SMA spinal cord (E-MTAB-3664^15^), human (GSE69175 ^67^, murine (GSE51735 ^68^) SMA motor neurons, and human spinal cord organoid (SCO; GSE290980 ^69^).

Pseudotime trajectory and correlation analyses were performed on log2-normalized data using Monocle 3 within Partek® Flow, with ventral or dorsal control clusters specified as the root. Pseudotime correlations were calculated using Pearson (linear) correlation. Gene-specific correlation analysis in the SMA ventral interneuron cluster was performed using median ratio–normalized data.

Spatial gene expression heatmaps and violin plots were generated with Partek® Flow using raw UMI expression unless otherwise stated. Expression heatmaps of gene families and categories were generated using ClustVis^70^ based on fold-change values within respective clusters. Hierarchical clustering was performed using average linkage and correlation distance without data scaling.

