## Supplemental Files for "Spatial transcriptomics identifies dysregulated programs across neural and non-neural tissues in spinal muscular atrophy"

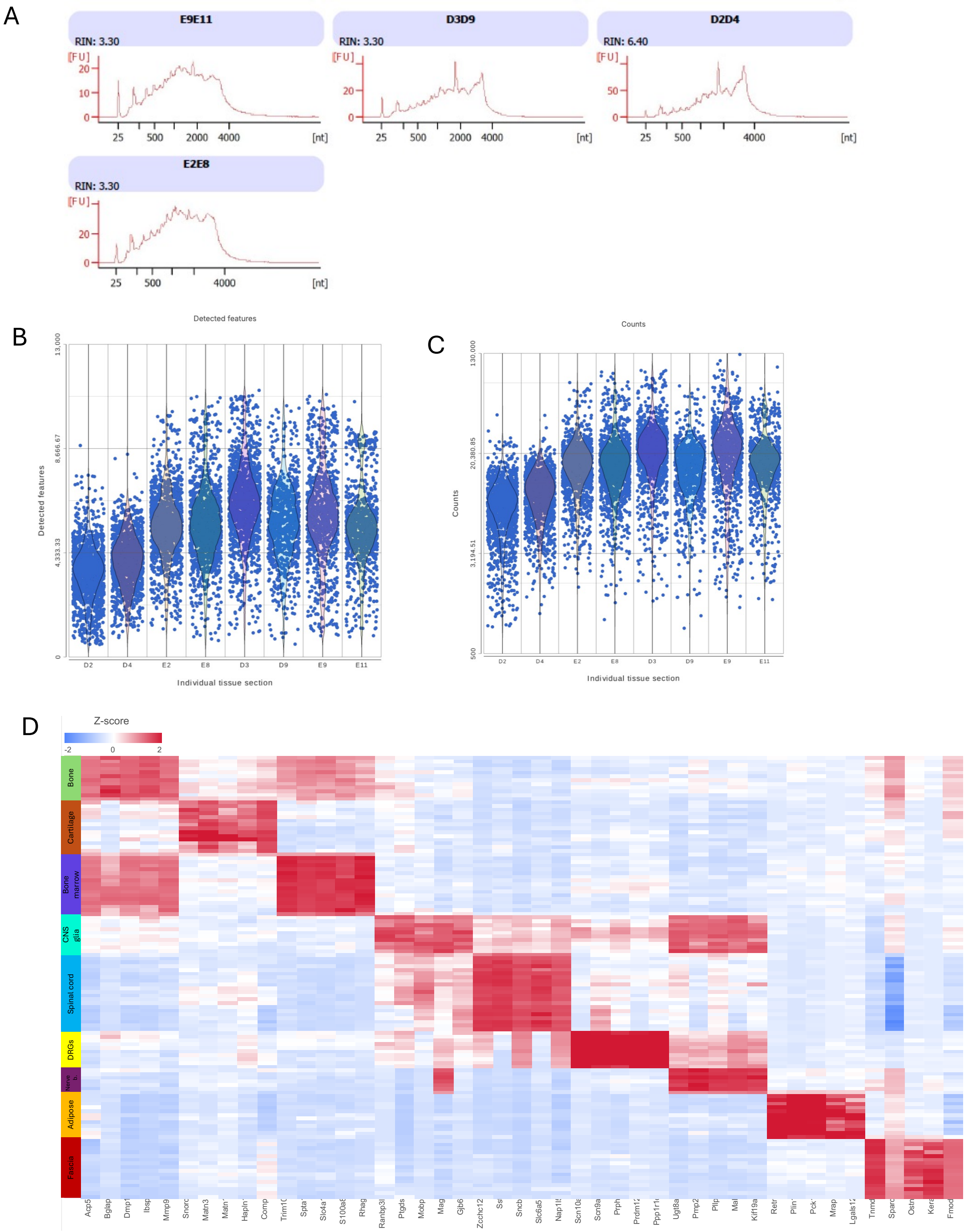


**Supplemental Figure 1.**

**A)** RNA integrity plots from formalin-fixed paraffin-embedded (FFPE) mouse tissue. DV200 was >93% for all samples. Heterozygous control animals (D2, D4, E2, E8) and SMA animals (D3, D9, E9, E11*)*. **B)** Violin plot (linear scale) illustrating the number of unique genes in each individual tissue section from heterozygous control animals (D2, D4, E2, E8) and SMA animals (D3, D9, E9, E11*)*. **C)** Violin plot (log scale) illustrating the number of unique molecular identifiers (UMIs) per spot in each individual tissue section from heterozygous control animals (D2, D4, E2, E8) and SMA animals (D3, D9, E9, E11*)*. D) Heatmap showing top 5 biomarker gene expression of cluster-defining genes across annotated non-muscle tissues.

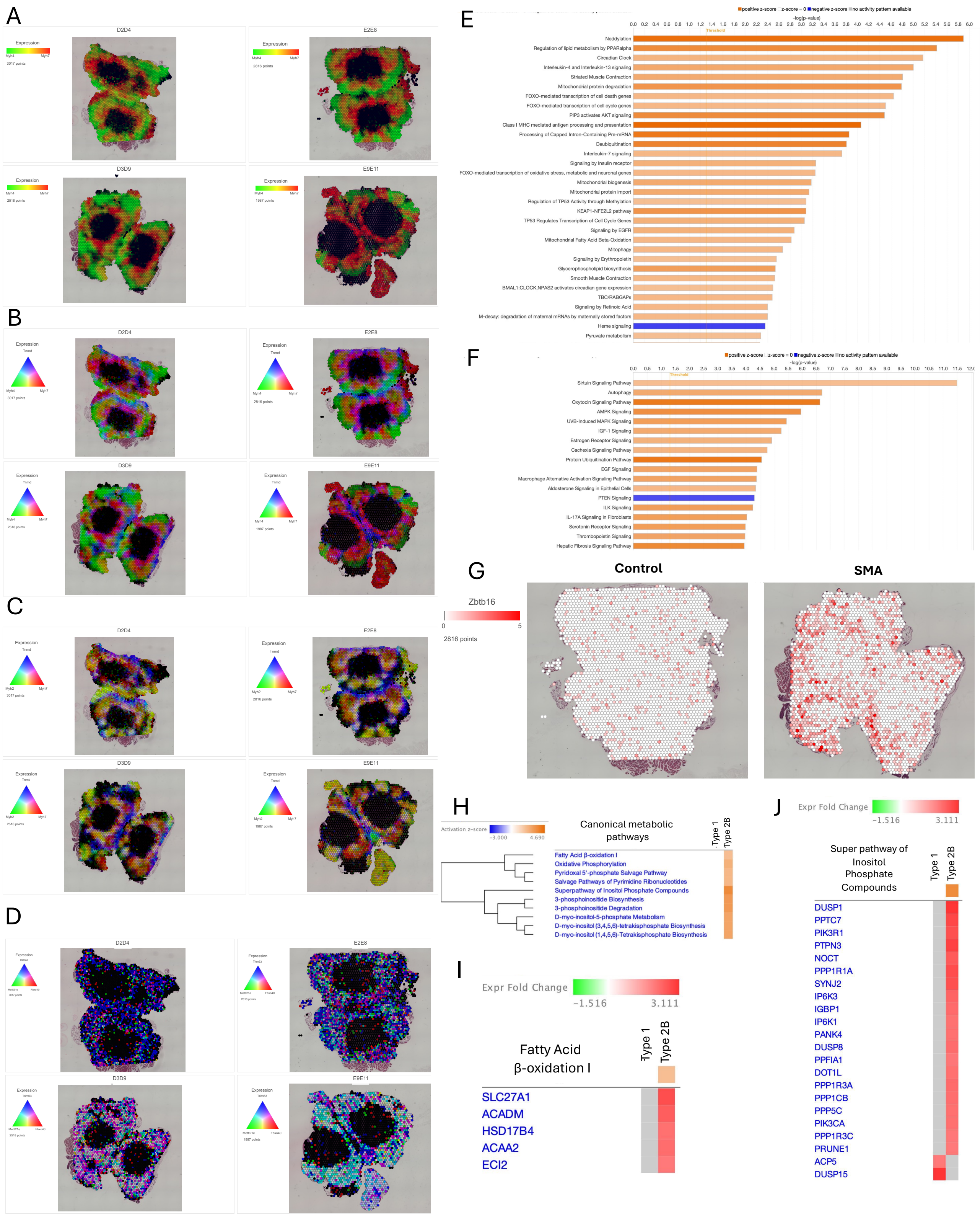


**Supplemental Figure 2.**

**(A)** Spatial heatmap of the spinal columns from control (D2D4, E2E8) and SMA animals (D3D9, E9E10) showing the log2 normalized mean expression of Type 1(*Myh7,* red) and Type 2b (*Myh4,* green) muscle markers. **(B)** Spatial heatmap showing gene marker expression for intramuscular connective tissue (*Tnmd*, blue) comparied to *Myh7* (red) and *Myh4* (green). **(C)** Spatial gene expression of Type 2a muscle (*Myh2*, blue) marker in comparison to *Tnmd* (blue) and *Myh7* (red). **(D)** Spatial gene expression of *Mettl21e* (green), *Fbxo40* (red), and *Trim63* (blue). Color intensity correlates with transcript abundance, and white spots indicate co-expression of all three genes. SMA Type 2b DEGs were analyzed with Qiagens IPA Reactome **(E)** or signaling **(F)** pathway analysis (z-score $\geq$2). **(G)** Spatial heatmap of raw UMI expression of *Zbtb16* in representative control and SMA animals. **(H)** SMA Type 2b DEGs were analyzed with Qiagens IPA metabolic pathway analysis (z-score $\geq$2). Gene expression heatmap of DEGs in **(I)** Fatty acid $\beta$-oxidation and **(J)** Super pathway of inositol phosphate compounds detected in SMA Type 2b muscle in comparison to SMA Type 1 gene expression.

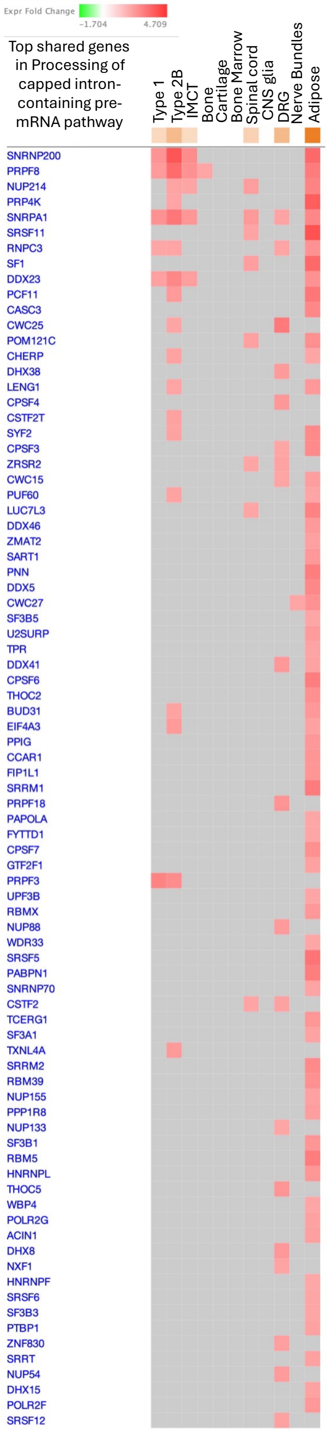


**Supplemental Figure 3.**

Extended Figure 2H showing all genes in cross-tissue comparison analysis of the top genes involved in the processing of capped intron-containing pre-mRNA pathway across multiple SMA tissues, including Type 1 and Type 2b muscle, bone marrow, cartilage, spinal cord, dorsal root ganglia (DRG), nerve bundles, and adipose tissue.

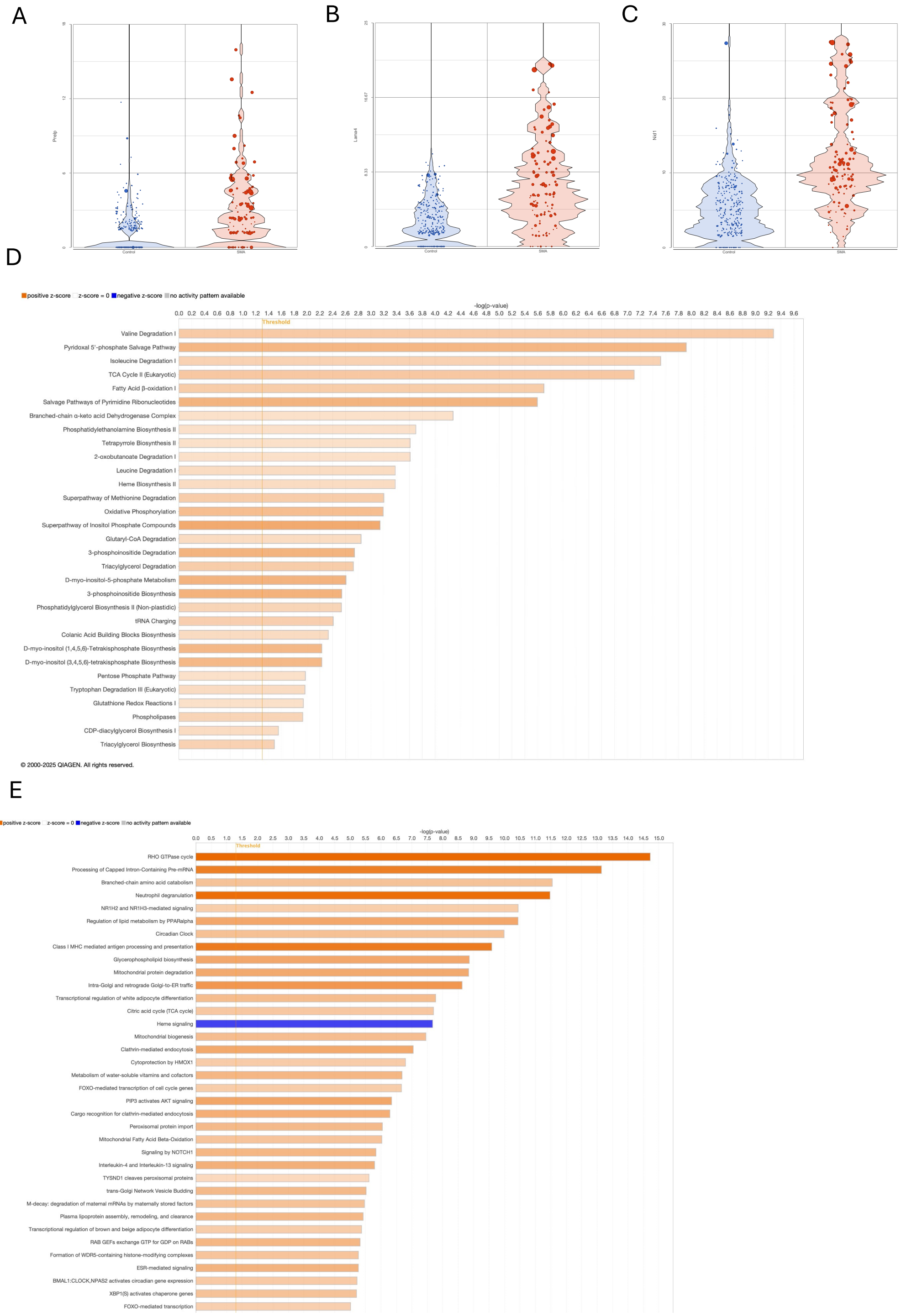


**Supplemental Figure 4.**

Violin plots showing spatial transcript levels of *Prelp* **(A)**, *Lama4* **(B)**, and *Nid1* **(C)** in control and SMA adipose tissue. Qiagens IPA metabolic **(D)** or reactome **(E)** pathway analysis (z-score≥2) of SMA adipose DEGs.

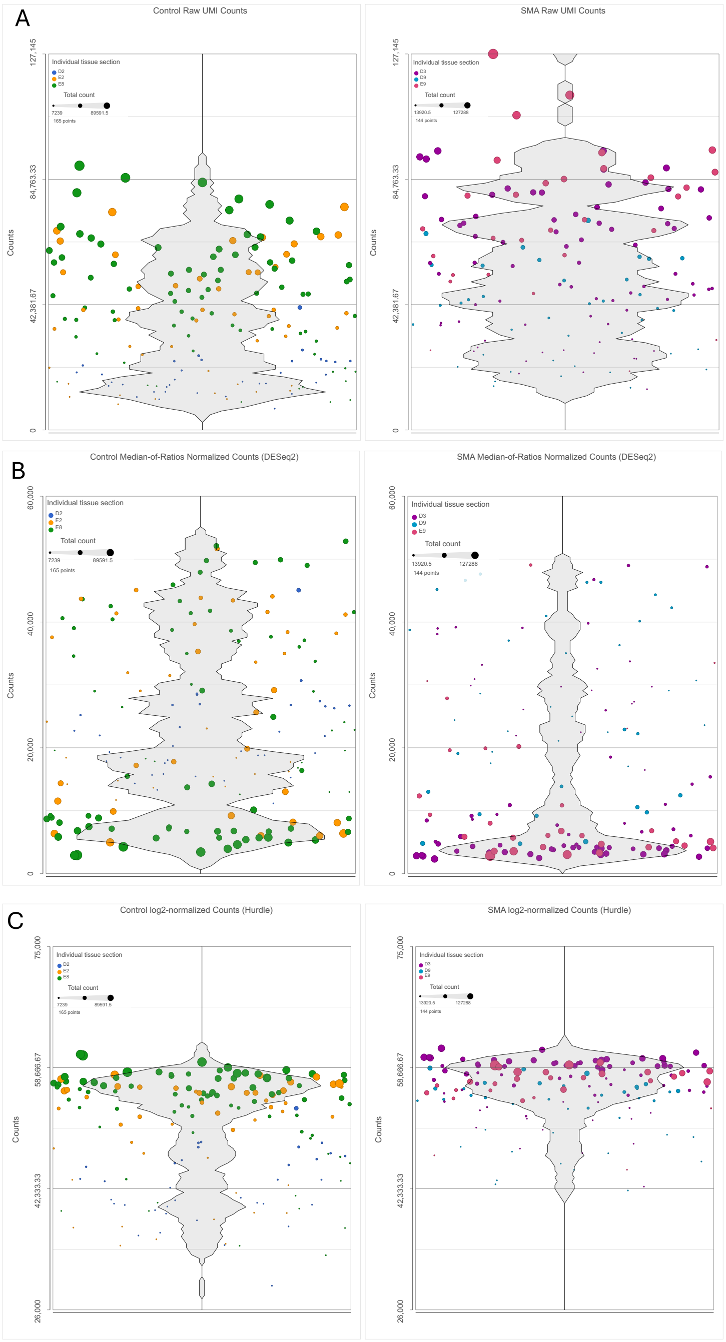


**Supplemental Figure 5.**

Raw UMI count (**A**), Median-of-ratios normalized count (DESeq2, **B**), and $log_2$-normalized count (**C**) distributions for ST spots in control (left) and SMA (right) DRG samples. Each point represents a ST spot, colored by individual animal ID with point size proportional to total counts.

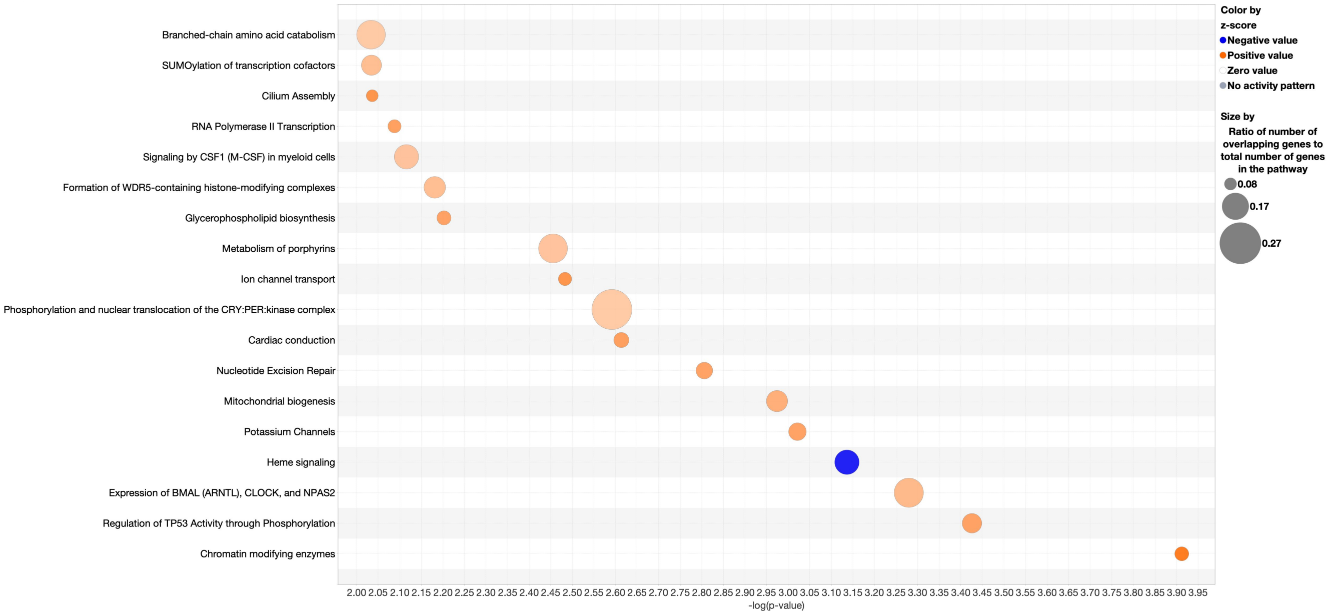


**Supplemental Figure 6.**

SMA DRG DEGs were analyzed with Qiagens IPA Reactome pathway analysis (z-score ≥2).

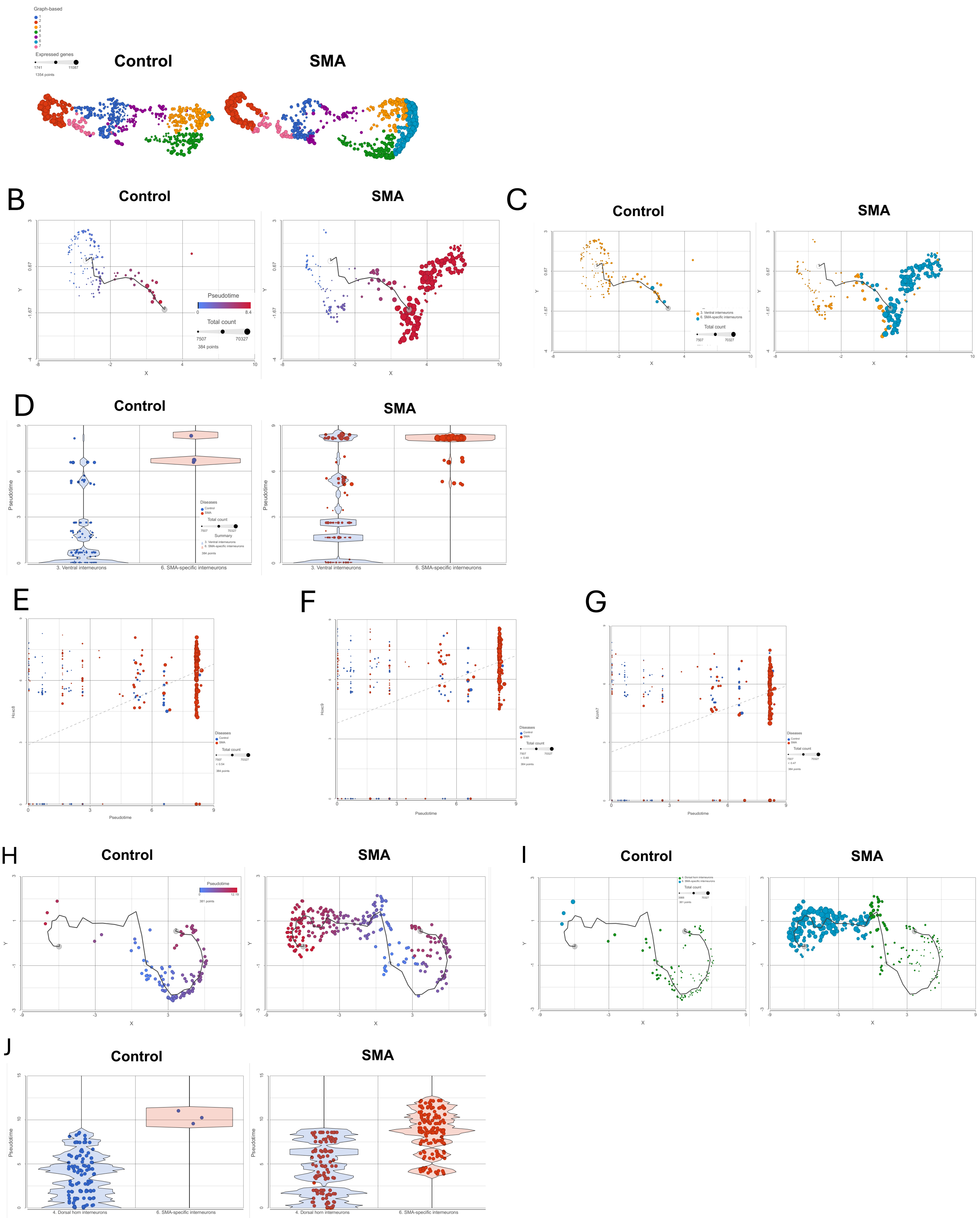


**Supplemental Figure 7.**

**(A)** UMAP representation of ST spots from control (left) and SMA (right) spinal cord sections, colored by graph-based clustering. Clusters corresponding to dorsal horn excitatory interneurons, ventral interneurons, and an SMA-specific cluster are preserved across conditions, with the SMA-specific cluster observed exclusively in SMA samples. ST size reflects number of expressed genes. Trajectory analysis

(Monocle 3) of ventral-derived ST spots performed for control (left) and SMA (right) samples **(B)** or graph-based clusters **(C)**, and the black curve denotes the inferred principal trajectory and size is adjusted for total UMI count. SMA-specific spots occupy late pseudotime states that extend beyond the range observed in control tissue, indicating a disease-exclusive transcriptional state with increased transcriptional activity. **(D)** Violin plots showing pseudotime distributions for selected ventral populations and the SMA specific cluster in control (left) and SMA (right) samples. Pseudotime values were calculated using a ventral-rooted trajectory, and individual points represent ST spots. SMA samples exhibit a significant shift toward higher pseudotime values, consistent with disease-associated progression to late transcriptional states. Correlation analysis of *Hoxc8* **(E)**, *Hoxc9* **(F)** and *Kcnh7* **(G)** gene expression along the ventrally rooted trajectory identified ST spots progressively increasing with pseudotime. Hoxc8 expression correlated with increased pseudotime (r=0.54), which was enriched with the SMA specific cluster (red). Trajectory analysis of dorsal-derived ST spots from control (left) and SMA (right) samples, with dorsal spots used as the root for pseudotime inference. ST spots are colored by pseudotime **(H)** or graph-based clusters **(I)**. **(J)** Violin plots showing pseudotime distributions for selected dorsal populations and the SMA specific cluster in control (left) and SMA (right) samples.

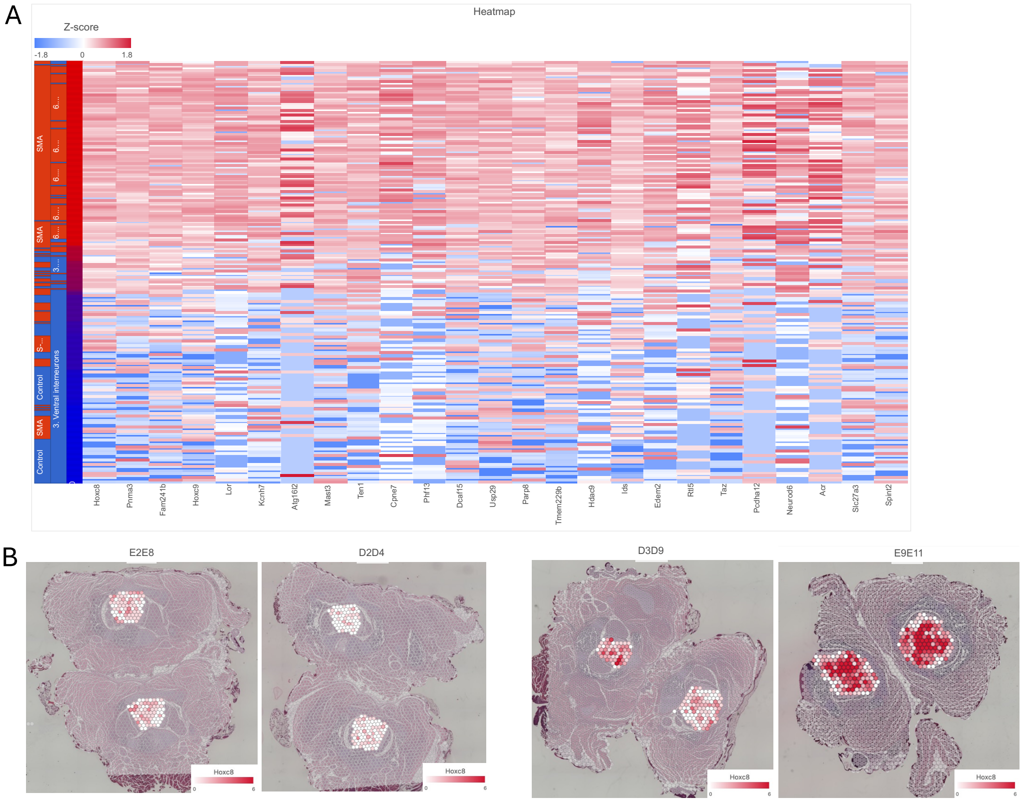


**Supplemental Figure 8.**

**(A)** Heatmap showing z-score normalized expression of genes that significantly correlate with pseudotime along the ventrally rooted trajectory. Columns represent genes and rows represent individual ST spots ordered by pseudotime. ST spots are annotated by condition and first- and second-order cluster identities (first order: Cluster 3, ventral interneurons; Cluster 6, SMA-specific cluster; second order: disease condition, SMA or control). Warmer colors indicate higher relative expression, whereas cooler colors indicate lower expression. **(B)** Spatial heatmap of *Hoxc8* gene expression in the spinal cord of control (E2E8, D2D4) and SMA (D3D9, E9E11) spinal cords.

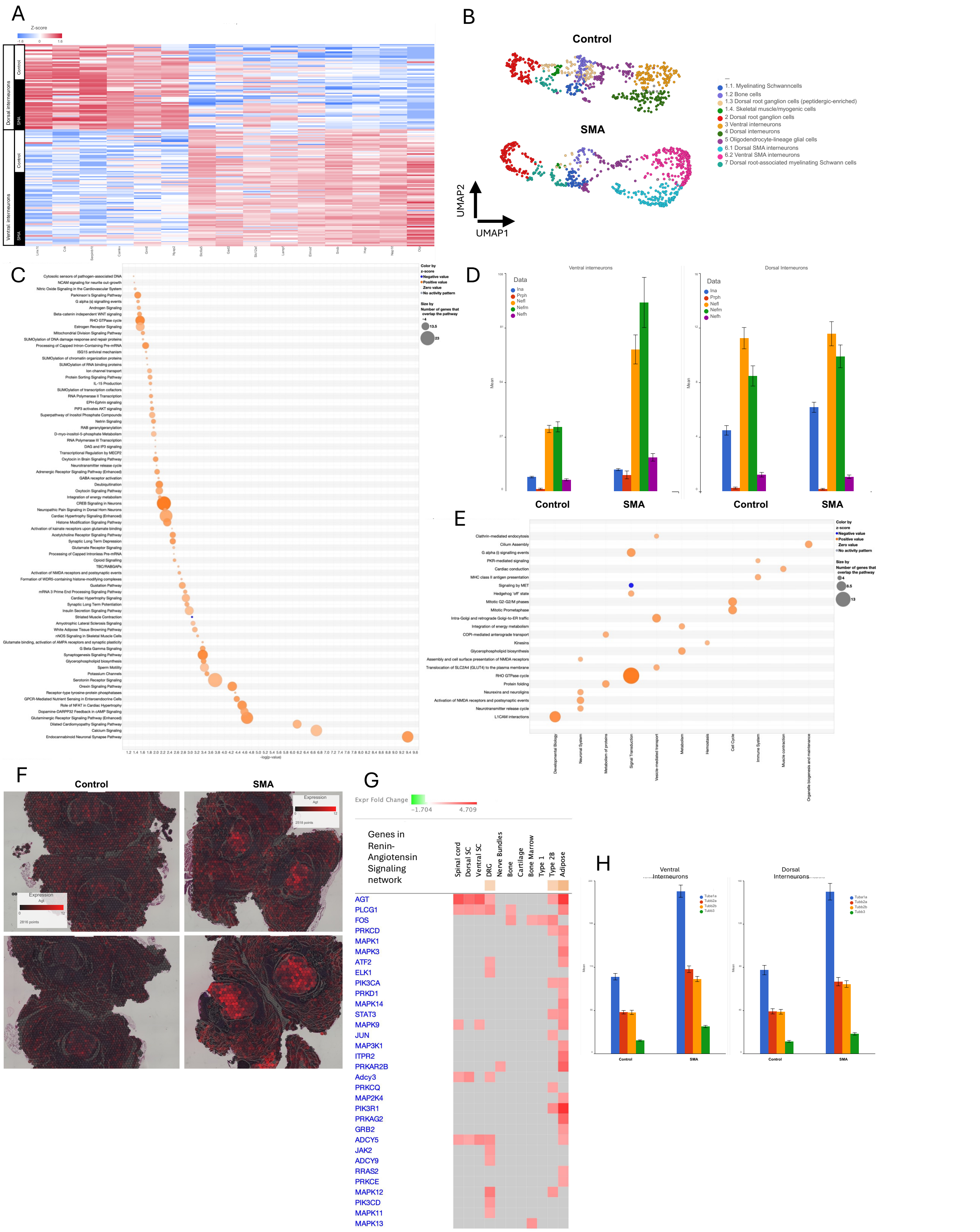


**Supplemental Figure 9.**

**(A)** Heatmap of z-score normalized expression of biomarker genes identified in the subclustering analysis of the SMA specific cluster (cluster 6). Rows represent ST spots ordered by ventral and dorsal interneurons annotated by condition (control or SMA). **(B)** UMAP visualization of ST spot clusters in control and SMA spinal cord samples showing that the sub clustering result of the SMA specific cluster. **(C)** IPA pathway enrichment analysis of DEGs identified in the dorsal region of SMA spinal cord. Dot size indicates the number of genes per pathway and color indicates enrichment direction. **(D)** Expression of selected genes across ventral and dorsal interneuron populations in control and SMA tissues. **(E)** IPA pathway enrichment analysis of DEGs identified in the ventral region of SMA spinal cord. **(F)** Spatial heatmap showing raw expression of *Agt* in control and SMA tissue sections. Increasing red intensity correlates with increasing *Agt* expression. **(G)** Gene expression heatmap of DEGs in the Renin-Angiotensin Signaling network across multiple SMA tissues. Increasing red intensity correlates with increasing gene expression. **(H)** Quantification of pathway-associated transcripts in control and SMA samples.

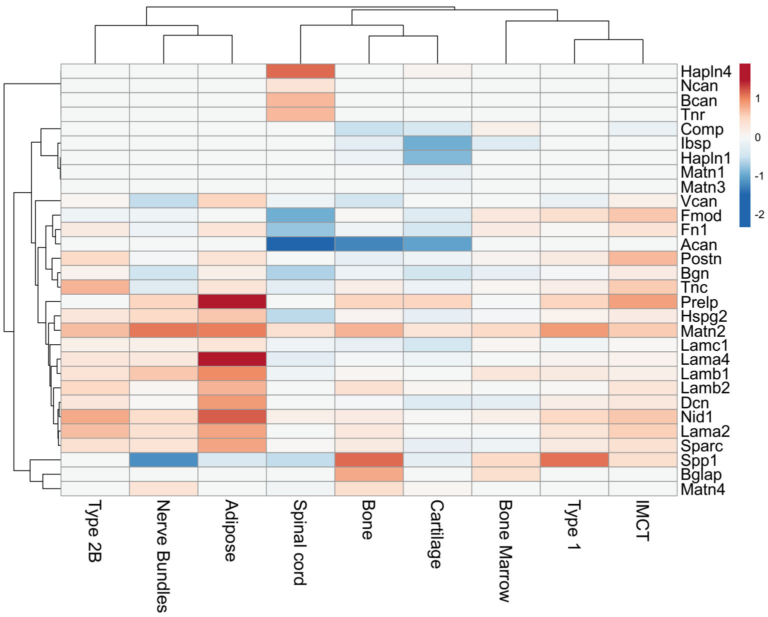


**Supplemental Figure 10.**

Hierarchical clustering heatmap of select ECM genes across annotated tissue regions. Color scale represents fold-change expression in SMA tissue.

### Supplemental Tables

**Supplemental Table 1.** Summary of experimental mouse phenotypes and Visium statistics.

**Supplemental Table 2.** Functional classification of the top 10 Biomarkers in Type1 SMA muscle.

**Supplemental Table 3.** Top 10 Biomarker genes identified in the tissue clusters identified in control and SMA spinal column cross-sections.

**Supplemental Table 4.** DEGs identified in the SMA type 2b muscle cluster.

**Supplemental Table 5.** DEGs identified in the SMA type 1 muscle cluster.

**Supplemental Table 6.** DEGs identified in the SMA bone cluster.

**Supplemental Table 7.** DEGs identified in the SMA cartilage cluster.

**Supplemental Table 8.** DEGs identified in the SMA bone marrow cluster.

**Supplemental Table 9.** IPA based upstream regulator prediction in SMA bone marrow.

**Supplemental Table 10.** DEGs identified in the SMA adipose cluster.

**Supplemental Table 11**. Selective DEGs in SMA adipose and their canonical biological functions.

**Supplemental Table 12.** DEGs identified in the SMA DRG cluster.

**Supplemental Table 13.** DEGs identified in the SMA dorsal interneuron cluster.

**Supplemental Table 14.** DEGs identified in the SMA ventral interneuron cluster.

**Supplemental Table 15.** Fold-changes of complement pathway members in select SMA tissues.

**Supplemental Table 16.** Fold-changes of collagen family members in select SMA tissues.

**Supplemental Table 17.** Fold-changes of ECM proteins mRNA expression in select SMA tissues.

**Supplemental Table 18.** Fold-changes of ankyrin family members in select SMA tissues.

| File name | Animal ID | Pheno-type | Sex | Number of spots under the Tissue | Mean reads per spot | Median Genes per Spot | Number of Reads | Genes detected | Reads Mapped Confidently to the Filtered Probe Set |
|---|---|---|---|---|---|---|---|---|---|
| D2D4 | D2 and D4 | Control | F | 3017 | 42,793 | 3786 | 129,105,917 | 19,272 | 88.80% |
| E2E8 | E2 and E8 | Control | F | 2,816 | 44,058 | 5,466 | 124,067,491 | 19,297 | 89.00% |
| E9E11 | E9 and E11 | SMA | F | 1,987 | 49,905 | 5,595 | 99,162,205 | 19,288 | 90.00% |
| D3D9 | D3 and D9 | SMA | F | 2,518 | 48,840 | 6,104 | 122,978,806 | 19,305 | 89.40% |
| **average** | | | | **2,585** | **46,399** | **5,238** | **118,828,605** | **19,291** | **89.30%** |
| **SEM** | | | | **224** | **1,750** | **503** | **6,689,921** | **7** | **0.26%** |

**Supplemental Table 1.** Summary of experimental mouse phenotypes and Visium statistics.

| Pathway | Genes | Biological Function | Effect In Disease |
|---|---|---|---|
| Mitochondrial/Metabolic Response | Nmrk2, Perm1, Retsat | $NAD^+$ salvage, mitochondrial biogenesis, lipid oxidation | Adaptive early, may become inefficient under sustained stress |
| Contractile Remodeling | Myh7b, Myl10 | Fiber-type switching, sarcomere reorganization | Adaptive remodeling that may become maladaptive with sustained stress |
| $Ca^{2+}$ & Thermogenic Stress | Sln | SERCA uncoupling, thermogenesis | Associated with increased energy expenditure and reduced contractile efficiency |
| Proteostasis / Regeneration | Fbxo40, Mettl21e | Protein turnover, myofiber regeneration | Reflects activation of protein turnover and stress-response pathways associated with atrophy and regeneration |
| Circadian & Metabolic Alignment | Per1 | Stress-induced circadian reprogramming | Links circadian regulation to metabolic and stress-response pathways |
| Electrical Remodeling | Kcnj12 | Membrane excitability | May alter membrane excitability and electrical stability |

**Supplemental Table 2.** Functional classification of the top 10 Biomarkers in Type1 SMA muscle.

| Upstream Regulator | Expr Fold Change | Molecule Type | Predicted Activation State | Activation Z-Score | P-Value of overlap | Target Molecules in Dataset |
|---|---|---|---|---|---|---|
| Lipopoly-saccharide | | chemical drug | activated | 4.138 | 8.99E-21 | *Alox5ap,Anxa1,Apobr,Ccl9,Ccn1,Cd33, Cd37,Cd52, Chi3l1,Crispld2,Csf3r,Cst3,Cxcr2,Dclre1c,Ddit3,Ddit4, Fkbp5,Fos,Fpr1,Fpr2,Fst,Fyb1,Hopx,Hp, Il1rn,Il36g,Jdp2, Kit,Lcn2,Mapk13, Mmp8,Mt1,Mt2a,Ncf1,Olfm4,Pag1, Pawr,Pglyrp1,Podxl,Ptx3,S100a8,Sell,Selplg,Slfn2,Stxbp2, Syk, Tap2,Tf,Tsc22d3,Vav1,Vcam1,Xdh,Zfp36l2* |
| TNF | -1.005 | cytokine | activated | 4.732 | 1.46E-16 | *Adam8,Alox5ap,Anxa1,B4galnt1,Ccl6, Ccl9,Ccn1,Cd33, Chi3l1,Crispld2,Cxcr2,Ddit3,Ddit4,Fkbp5,Fos,Fpr1,Fpr2, Fst,Hopx,Hp,Il1rn,Il36g,Kit,Lcn2,Mmp8,Mt1,Mt2a,Ncf1, Pglyrp1,Podxl,Ptx3, S100a8,Sell,Selplg,Slfn2,Syk,Tap2, Tf, Tsc22d3,Vav1,Vcam1,Xdh* |
| IL1B | 1.071 | cytokine | activated | 4.696 | 4.09E-16 | *Adam8,Anxa1,Ccl9,Cd177,Chi3l1, Crispld2,Ddit3,Ddit4, Fkbp5,Fos, Fpr2,Fst,Gas6,Hopx,Hp,Il1rn, Il36g,Lcn2, Lcp1,Mapk13,Mmp8, Mt1,Mt2a,Olfm4,Podxl,Ptx3, S100a8,Slfn2,Tap2,Tsc22d3,Vcam1,Xdh* |
| TCL1A | 1.001 | transcription regulator | | 1.3 | 1.10E-13 | *Csf3r,Cxcr2,Fkbp5,Fos,Fpr1,Lcn2,Mmp8,Mpeg1,Ngp, Retnlg,S100a8* |
| CEBPE | 1.394 | transcription regulator | activated | 2.613 | 1.11E-13 | *Alox5ap,Ccl9,Csf3r,Il1rn,Lcn2,Mmp8,Ncf1,Ngp,Retnlg, Sell* |

**Supplemental Table 9.** IPA based upstream regulator prediction in SMA bone marrow.

| Function | Maladaptive Genes | Compensatory Genes |
|---|---|---|
| Rhoa/rock activation and cytoskeletal contractility | Arhgef1, Arhgef17, Net1, Pkn2, Diaph2 | Arhgap12, Arhgap21, Arhgap35, Dlc1, Myo9b, Ophn1 |
| ECM mechanosensing and fibrotic differentiation | Itgb1, Vim | Rhou, Fam13a |
| Growth-factor/rtk pro-fibrotic signaling | Grb2, Frs2 | |
| Ros generation and oxidative stress | Cyba | Hint2 |
| Mitochondrial quality control and protection | | Samm50, Hint2 |
| Cytoskeletal remodeling and stress-fiber turnover | Diaph2, Vim | Ophn1, Myo9b, Rhou |
| Metabolic adaptation and lipolysis–mitochondrial crosstalk | | Fam13a, Samm50, Hint2 |

**Supplemental Table 11.** Selective DEGs in SMA adipose and their canonical biological functions.
